# Self-Assembly of a Repeatable DNA Nanohinge System Supporting Higher Order Structure Formation

**DOI:** 10.1101/2023.05.26.542516

**Authors:** Melanie Law, Charlie Susham, David Mackay, Stephanie Nguyen, Rachael Nicholas, Miguel Roberto G. Tsai, Ethan Rajkumar, Fumiya Inaba, Kieran Maheden, Immanuel Abdi, Joe C.H. Ho, Brandon Kieft, Steven J. Hallam

## Abstract

DNA base pairs can both encode biological information and be used as a programmable material to build nanostructures with potential application in nanofabrication, data processing and storage, biosensing and drug delivery. Over several decades development of these DNA origami nanostructures has led to increasingly advanced self-assembling nanostructures and molecular machines actuated by various mechanisms such as toehold-mediated strand displacement (TMSD), magnetism and even light. However, scalability remains challenging as using larger scaffold strands can increase the likelihood of kinetic traps and misfolded conformations. Here we describe a repeatable DNA nanohinge system to increase the scalability of existing nanohinge designs for hierarchical assembly of more complex structures with greater degrees of mobility and functionality. The components of this system, comprising two distinct nanohinges, were designed in caDNAno. Structure conformation and stability were simulated using CanDo and MrDNA, and hinge assembly was validated by TEM. Electron micrographs revealed hinge-shaped nanostructures capable of self-assembly into more complex structures, as well as actuation using TMSD through a reversible locking mechanism incorporated into the design. Our work expands the existing utility of DNA nanohinges as building blocks for scalable DNA nanostructures and demonstrates the feasibility of polymerizing hinges in a novel manner for higher order assembly. The enhanced functionality of our dual hinge systems can be employed in future applications requiring greater control and mobility of DNA nanostructures.

## INTRODUCTION

Programmable DNA origami can be used to construct self-assembling nanostructures and molecular machines including walkers, tweezers, and boxes, that can be actuated in response to intrinsic or extrinsic signals to specify a structural change or defined movement [1]. Potential application of DNA origami in nanofabrication, data processing and storage, biosensing and drug delivery have motivated development of increasingly complex designs [2] actuated using a number of different mechanisms including temperature [3], pH [4], toehold-mediated strand displacement (TMSD) [5], magnetism [6], and even light [7]. For example, Lauback and colleagues developed DNA origami rotors and hinges responsive to magnetic actuation to direct self-assembly in real time [6]. Kuzyk and colleagues developed a DNA origami nanostructure responsive to light-driven amplitude modulation paving the way for more advanced optical devices and sensing platforms [7].

Using the technique of DNA origami, DNA nanostructures can self-assemble through complementary base pairing between short oligonucleotide ‘staples’, and a longer single-stranded (ss) DNA ‘scaffold’, typically between 7-8 kilobase pairs long. The specific binding of the staples to the scaffold causes the scaffold to fold into a programmed 2D or 3D conformation [8]–[10]. Scaffold-staple base pairing layouts are currently designed using software applications such as caDNAno [11] and optimized using stability and structure prediction algorithms implemented in CanDo [12] and MrDNA [13], respectively. DNA nanostructure size can be increased by using a longer scaffold strand with more staples [14], [15]. However, this introduces design challenges by increasing the possibility of kinetic traps and misfolded conformations [16], [17]. Hierarchical DNA assembly of smaller DNA nanostructures with less staples represents a way to scale-up structure size, while minimizing scaffold–staple crossover density. Examples of foundational units for higher order DNA structures include DNA tiles [18]–[20], bricks and hinges [6], [21], [22], and tripods [23]. Specifically, the unique characteristics of DNA origami hinges make them versatile building blocks in the field of nanotechnology, enabling the construction of sophisticated and functional nanoscale devices, drug delivery systems, sensors, and other applications that require controlled motion and precise structural arrangements.

DNA nanohinges typically consist of two stiff bundles (bricks) of double stranded (ds) DNA helices connected by ssDNA loops of varying lengths that can open and close with a single degree of rotational freedom not unlike a macroscopic hinge [5], [6]. These bricks constitute the arms of the hinge. Nanohinge functionality is realized through opening, closing or locking at a designated angle [24], which has been achieved for nanohinges and hinge-like structures by adjusting the ssDNA connection length [25], attaching magnetic beads [6], and using TMSD [26]. These mechanisms have enabled hinges to open a nanoscale box [27], investigate DNA unwrapping from a nucleosome [28], [29], and localize dyes with Bohr-radius resolution [30].

Although nanohinges represent an important component in assembling higher-order structures for nanotechnology, a scalable and actuatable DNA nanohinge system assembled from single hinge units has yet to be developed. Scalability of DNA nanostructures presents a major challenge in enabling the practical use of nanomaterials [31].

Here we describe a repeatable DNA nanohinge system intended to circumvent certain design limitations related to higher order nanostructure formation. Our nanohinges are composed of a single scaffold strand which folds into two dsDNA nanobricks connected by seven ssDNA loops forming the pivot point of the hinge with a TMSD actuated locking mechanism incorporated into the design. Due to the directionality of each scaffold helix, two hinges denoted as H1 and H2 were designed with the pivot point on different edges. As a result, H1 and H2 can polymerize in an alternating sequence to form higher order nanohinges (HONs). TMSD is an integral mechanism used in many DNA origami nanostructures to control interaction kinetics based on the process of branch migration between complementary DNA strands [32]. With respect to our design, TMSD allowed single hinges to be locked at predetermined angular conformations depending on the presence of specific oligonucleotide sequences. This reversible locking mechanism in combination with hierarchical assembly of single hinge units into more complex structures provides both increased degrees of movement and enhanced functionality with potential nanofabrication and biosensing applications.

## METHODS

### DNA Nanohinge Design and Simulation

DNA nanohinge system components were designed in caDNAno2 [11] using the p8064 scaffold. H1 and H2 were designed with 56 helices, forming two 28-helix bricks for each hinge, connected in a honeycomb lattice. The free ends of H1 and H2 were left single stranded to accommodate polymerization strand binding for hinge-to-hinge polymerization. Polymerization strand sequences were manually determined from the H1 and H2 caDNAno2 files. Overhang sequences of the padlock strands were determined by selecting staple strands located at the peripheral helices of the H1 and H2 bricks. The remaining padlock and four other locking-mechanism strand sequences (lock, key, antikey, and antilock) were manually determined to prevent undesirable secondary structure formation as determined by NUPACK web application (Version 4.0 for local).

CaDNAno2 files generated for each hinge were processed in the CanDo web application (Version 1.1) to determine their three-dimensional conformation and relative regional flexibility. Structure stability was visualized using heatmaps with shading from blue to red, indicating areas of low to high relative root mean square fluctuation (RMSF) values, respectively, in that region. Default values for average B-form DNA geometry and DNA mechanical properties were used in CanDo. The output .bild file provided by the CanDo web application was viewed using UCSF Chimera (Version 1.16) [33], which rendered the structure in a three-dimensional interface, allowing for more detailed examination of the computed RMSF map on the structure. H1 and H2 structures in their equilibrium state were predicted using MrDNA (Version 0.2.0), a multi-resolution structure prediction software developed by Christopher Maffeo and the Aksimentiev group at the University of Illinois [13], and visualized using Visual Molecular Dynamics (VMD), an open-source software distributed by the Theoretical and Computational Biophysics group at the University of Illinois.

### DNA Nanohinge Optimization

In addition to holistic structural simulations to model structure stability, the staple strands of each hinge were analyzed for the presence of sandwich strands and kinetic traps. Staple strand analysis was performed using MATLAB (Version 2020b) and Python 3.9. Sandwich strands and kinetic traps identified by the scripts were removed in caDNAno2 to optimize structure stability and decrease the probability of structure misfolding. These scripts can be found at https://github.com/ubcbiomod/Higher-Order-Nanohinge-Systems. The caDNAno designs were further adjusted if the observed CanDo model displayed a red color in regions where it was not expected, as lower stability in unexpected regions may decrease the probability of proper structure folding and retention. CaDNAno designs were adjusted by altering the staple strand connectivity within a region of interest by either increasing or decreasing the number of crossovers between helices.

### H1 and H2 Hinge Assembly

To form H1 and H2 single hinge units, 20 nM p8064 (Tilibit nanosystems) scaffold was mixed with staple strands at a 1 to 5 molar ratio with 1x folding buffer (1 mM EDTA, 5 mM NaCl, 5 mM Tris), and 20 mM MgCl_2_. The staple strand master mix contained each staple required for hinge formation in a 1:1 molar ratio at 100 nM. Assembly was performed under a 37-hour thermal annealing ramp, adapted from Douglas et al. [7]. Assembly in 20 mM MgCl_2_ was undertaken to promote assembly and reduce smearing during gel analysis (Supplementary Fig. 6). Folding reactions were heated to 80°C, cooled to 60°C at a rate of 1°C per hour, then cooled to 24°C at a rate of 1°C per minute, then cooled to 4°C.

H1 and H2 assembly was determined by a 1% agarose gel in 1X TAE with 11 mM MgCl_2_. Running buffer was supplemented with the same conditions. Single hinge units were purified by spin column purification performed with 100 kDa Amicon Ultra Centrifugation filters (MilliporeSigma). The filters were equilibrated by centrifuging 500 μL of resuspension buffer (0.5X TAE, 4 mM MgCl_2_) at 5000 rcf for 15 minutes. The flow-through was discarded and an unpurified hinge solution was added to the column. The column was filled to 500 μL with resuspension buffer and centrifuged at 2000 rcf for 30 minutes. A wash step with resuspension buffer was included at the same centrifugation conditions as the previous step. The filter was inverted in a new tube and centrifuged at 1000 rcf for 2 minutes. The expected volume of the purified hinge solution ranged from 20 to 25 μL. H1 and H2 purification was assessed by an agarose gel with the same conditions as their assembly. To form hinges capable of actuation, padlock strands were added during the assembly stage at the same concentration as staple strands and subjected to the same conditions.

### TEM Imaging

Transmission electron microscopy (TEM) was used to verify the proper assembly of DNA nanostructures. Samples were prepared for imaging by two different methods. First, the samples were prepared by depositing 3 to 5 μL of a sample at 0.01% (w/v) onto a glow-discharged CU 400 formvar/carbon coated grid. Grids were allowed to stand for 3 minutes prior to wicking the moisture away using a paper point and negatively stained with 10 μL of saturated uranyl acetate (UA). Alternatively, the samples were prepared by mixing 5 μL of a sample with 5 μL of saturated UA; all 10 μL were then deposited on the same grid surface as the first method and were allowed to stand for 3 minutes. The remaining moisture was wicked away followed by a second application of UA. These two methods were applied interchangeably on a sample-by-sample basis based on the advice of a technician at the UBC Bioimaging Facility (BIF). Grids were imaged at either 80 kV (Hitachi H7600) or 200 kV (FEI/Thermo Tecnai G20). Imaging was performed by the UBC BIF.

### Locking Mechanism Verification

The locking mechanism verification was done in two parts with the first investigating the locking mechanism strand binding patterns, and the second investigating the effect of the locking mechanism on purified H1 and H2. The locking mechanism strand binding patterns were investigated to verify that all the strand combinations in the mechanism were able to bind as expected. The conditions which all included the padlock strand were as follows: lock, lock + antilock, lock + key, lock + key + antikey, lock + antilock + key + antikey, and key were incubated at 30 °C for 25 minutes. The strands were incubated in a 1:1 molar ratio, 12.5 mM MgCl_2_. Each locking mechanism strand was used individually as a negative control. All conditions were run on a 20% polyacrylamide gel for 2 hours at 115 V. Next the ability of the antilock and key strand to displace the lock strand from the padlock strand as well as the ability of the antikey to displace the key strand bound to padlock were evaluated. Padlock was incubated with lock or key in a 1:1 molar ratio, 12.5 mM MgCl_2_ at 30 °C for 25 minutes. Antilock was then added to padlock + lock, key added to padlock + lock, and antikey added to key + padlock and incubated as above. All conditions were run on Sodium Dodecyl Sulphate-Polyacrylamide Gel Electrophoresis (SDS-PAGE) as above. All conditions were repeated for each padlock strand of H1 and H2. To demonstrate hinge actuation via the locking mechanism, key or lock strands were incubated with H1 and H2, assembled with their respective padlock strands, at 2:1 molar ratio and incubated at 30 °C for 25 minutes. Analysis of TEM images to determine differences in hinge conformation states were performed in R Statistical Software (Version 4.1.2) [34]. The following R packages were also employed: tidyverse (Version 1.3.1), ggpubr (Version 0.4.0), rstatix (Version 0.7.0) and dyplyr (Version 2.1.1) [35]–[38].

### Higher Order Nanohinge Assembly

HONs composed of >1 single hinge pairs were assembled after purifying single H1 and H2 hinge units. H1 and H2 hinge units were connected via polymerization and neighbor strands in a 1.25:1 molar ratio. Sequences for polymerization strands are disclosed in Supplementary Table 3. Neighbor strands were excluded in single hinge assembly to prevent clumping of hinge units and were included during assembly to increase edge stability for hinge polymerization. Sequences for neighbor strands are disclosed in Supplementary Tables 2A and 2B. HONs were formed by incubating purified H1 and H2 at a final concentration of 5 nM each with polymerization and neighbor strands at a 7.5 and 6 molar excess, respectively, in ddH_2_O with 20 mM MgCl_2_. Samples were incubated at 45^σ^C decreasing by 2°C every hour until reaching 4^σ^C, adapted from Lauback et al. [6]. Successful assembly was verified on an agarose gel followed by TEM using the same method implemented for individual DNA nanohinges.

## RESULTS

### Structure Design of DNA Nanohinge Components

DNA nanohinge system components were designed to connect end-to-end, allowing for repeated polymerization of single hinge pairs (denoted as H1 and H2). H1 and H2 were constructed with two 28 helix cylindrical bricks connected by a single scaffold that created the hinges’ pivot point. H1 and H2 dimensions (length × width × height) were expected to be approximately 49.6 × 14.7 × 25.2 nm and 50 × 14.7 × 25.2 nm, respectively. These estimates were determined by multiplying the number of base pairs by 0.34 nm or the number of helical domain cross-sections by 2.1 nm along the appropriate plane of the hinge [10]. Consistent directionality of the scaffold ensured that only one end of an H1 brick connected to the complementary brick of H2 (Fig. 2). Incorporation of 30 partially complementary ssDNA strands to the scaffold of each H1 and H2 pair allowed for H1 and H2 polymerization into higher order structures. These polymerization strands were designed to have a minimum 7 bp overlap with the scaffold of connecting H1 and H2 helices. Scaffold-staple base pairing layout for the DNA nanohinge system is shown in Supplementary Fig. 1 and the staple sequences are provided in Supplementary Tables 2A and 2B, respectively.

**Figure 1.**
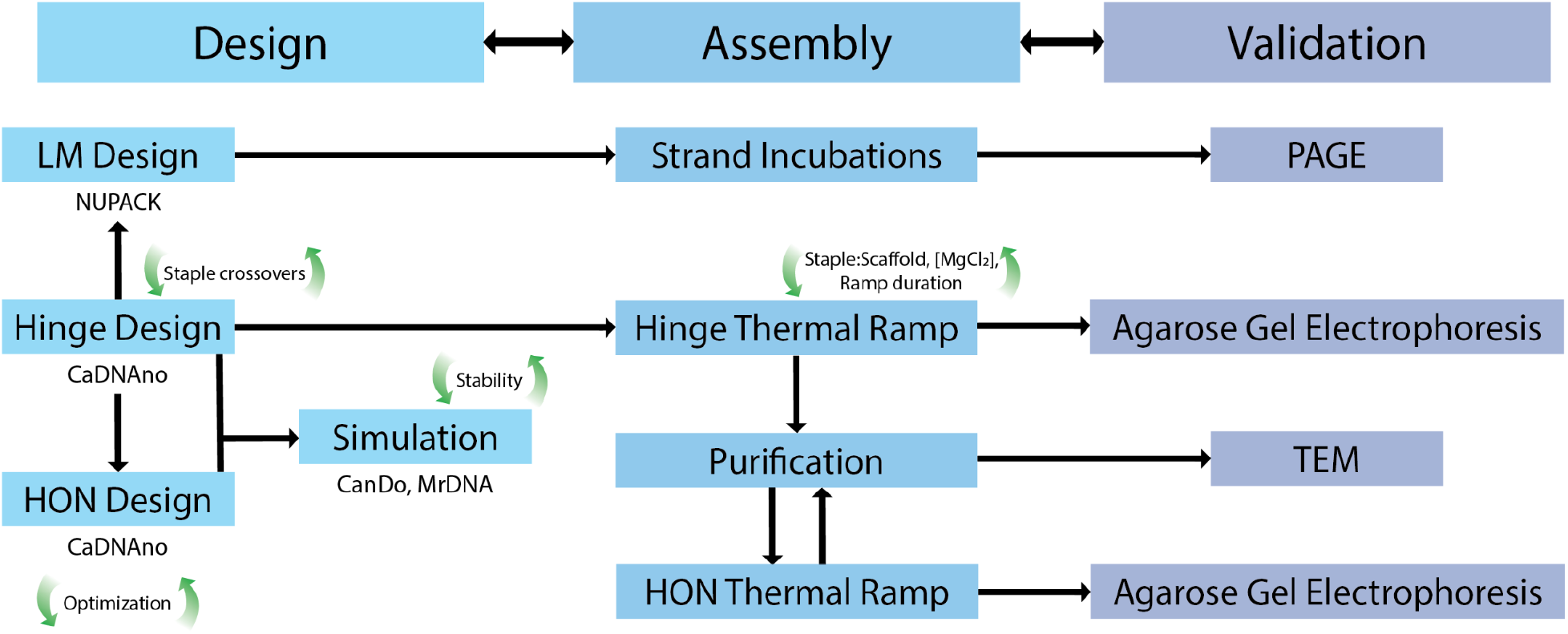
Workflow of the DNA nanohinge design, assembly and validation. The nanohinge, locking mechanism, and configuration for HONs were designed and simulated using software (CaDNAno, CanDo, MrDNA, NUPACK). The locking mechanism was assembled by incubation of varying strand combinations. Nanohinges were assembled by thermal ramps and purified by ultracentrifugation. Validation was performed using polyacrylamide gel electrophoresis (PAGE), agarose gel electrophoresis and TEM. LM = locking mechanism.

**Figure 2.**
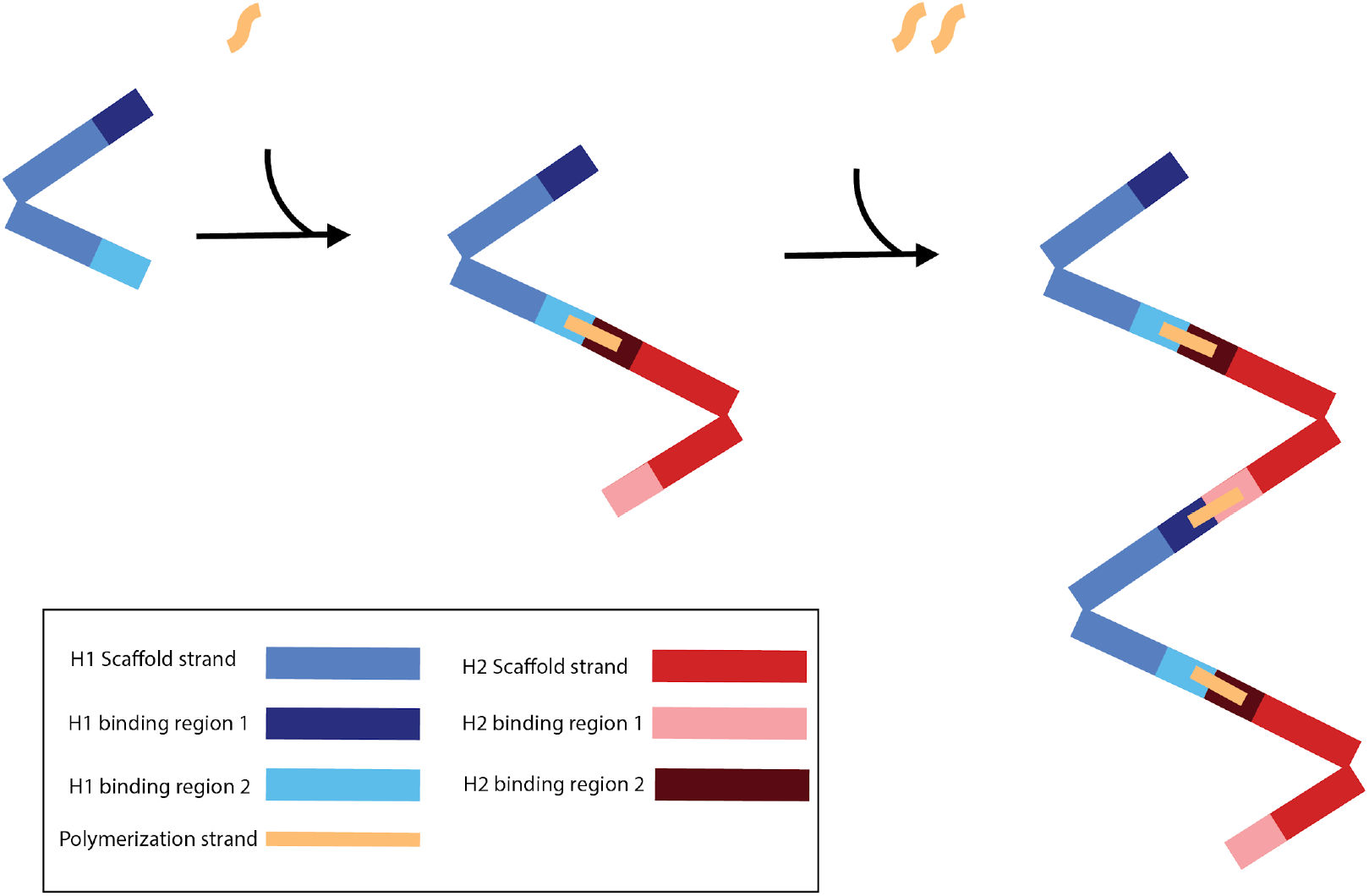
Schematic of the binding and the individual components of the DNA nanohinge system. Binding regions are color-coded to indicate binding directionality. H1 binding region 1 is uniquely complementary to H2 binding region 1, whereas H1 binding region 2 is uniquely complementary to H2 binding region 2. Polymerization strands are used to attach H1 and H2 together at the respective corresponding binding regions.

The design of each hinge was done using a single scaffold strand which crossed back and forth from the top brick and bottom brick, creating two distinct yet connected bricks. With respect to the staple strand connectivity, forced crossovers (i.e., crossovers which occur outside of their naturally occurring area as defined by the helical rotation of adjacent DNA helices) were largely avoided. Forced crossovers introduce structural tension by interfering with the standard helical rotation of B-form DNA [39], [40]. Despite this tension, it was necessary to introduce a few forced crossovers in consideration of overall structure stability [39]. For instance, at the pivot points of the H1 and H2 models, forced crossovers were implemented to ensure the structure did not fray from the single stranded pivot region strands and overly expose single stranded DNA regions. Because the region at the pivot point of the hinge was single stranded and the nucleotides are free to rotate, forced crossovers were not expected to introduce the same amount of structural tension as if it were introduced in fully double stranded B-form DNA.

Different versions of each structure were designed, built, and tested in an iterative process. For example, earlier iterations of H1 and H2, denoted as H1’ and H2’, were designed to connect end-to-end in a series of hinges along a single axis. Both H1’ and H2’ consisted of two 36 double helix bundles that were connected at one end by nine and eleven ssDNA staples, respectively. These staples were used to form the pivot point of the hinge (Supplementary Fig. 4). The expected dimensions of H1’ and H2’ were 43.5 × 20.3 × 13.5 nm and 42.8 × 20.3 × 13.5 nm, respectively. TEM images indicated that H1’ and H2’ hinges did not assemble properly due to a lack of observable hinge-like structures (Supplementary Fig. 5).

To render the hinge dynamic, an actuator mechanism was integrated based on the toe-hold mediated strand displacement (TMSD) mechanism previously used [32]. For this feature, a locking strand mechanism was incorporated to actuate H1 and H2. The mechanism consisted of six different ssDNA strands (Fig. 3). The padlock strand included overhangs that were complementary to the scaffold of each brick of a hinge. H1 and H2 both had two padlock strands bound to each side of the hinge. Complementary regions of the padlock binding to the locking strand forced the hinge to adopt a closed conformation. The unbound portion of the padlock was recognized by the key strand which bound and displaced the lock strand, opening the hinge into its extended conformation. The keystrand was displaced by the recognition of its overhang by the antikey strand, resetting the locking mechanisms. Bound lock strands were similarly displaced through hybridization with the antilock strand. Locking mechanisms sequences can be found in Supplementary Tables 4-5.

**Figure 3.**
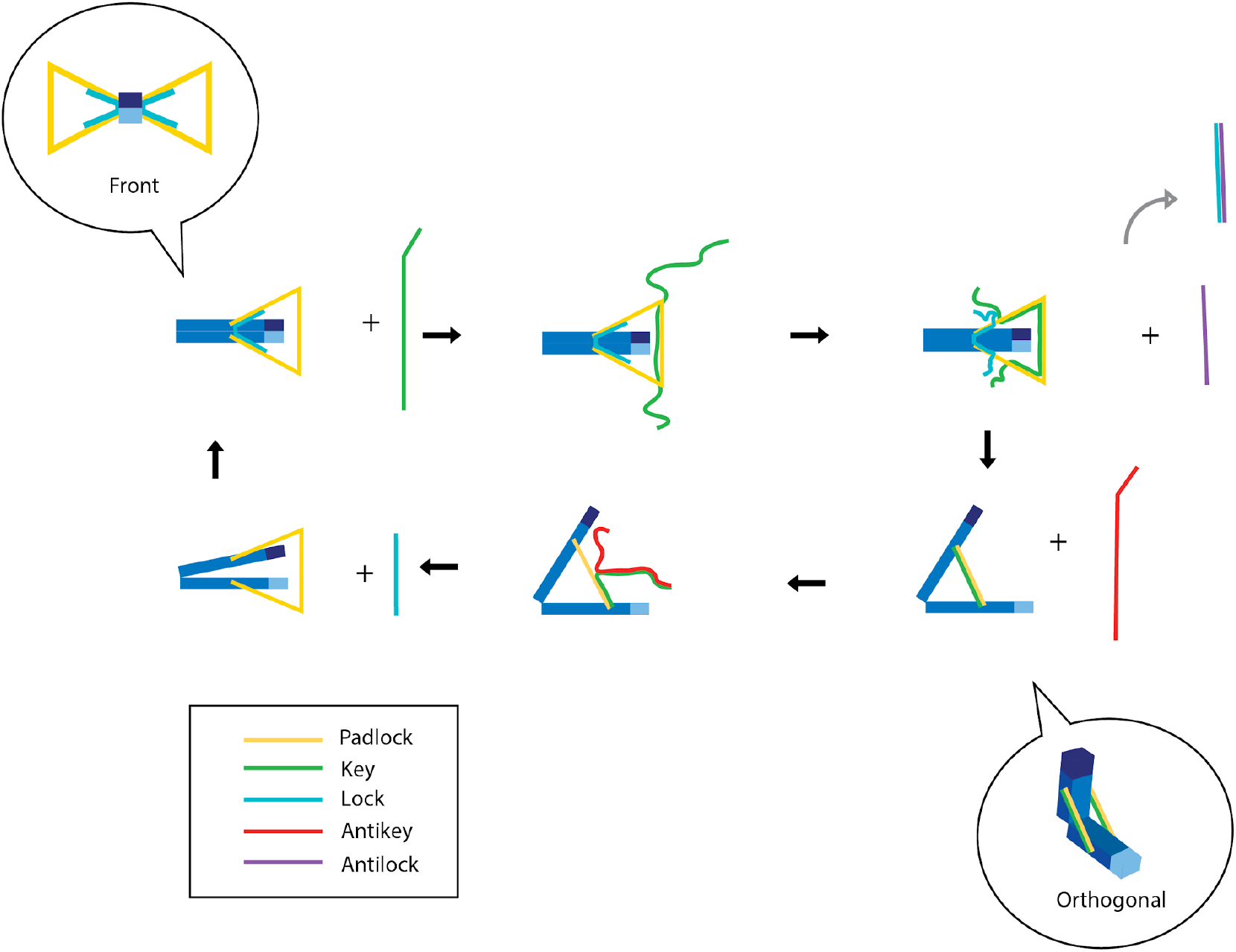
Locking Mechanism. The locking mechanism sequence depicted as a step-by-step process, in which complementary strands (lock, key, antilock, and antikey) are progressively added to replace an existing strand, thereby enabling the hinge to be opened or closed.

### H1 and H2 Modelling

Structure formation simulations were performed to estimate the integrity of H1 and H2 nanostructures at 25 °C using simulation software MrDNA [12]. The free ends of the hinge bricks were expected to have higher average root-mean-square deviation (RMSD) values, as nucleotides at the end of the hinge are more sensitive to angular movement [40]. Additionally, the ends of the hinge were left single stranded, fluctuating freer than if they were double stranded [12]. The use of the auto stapling feature in caDNAno produced structures with intermediate flexibility, which was indicated by their degree of shading based on RMSF values. Iterations of hinge designs with less high-flexibility areas in the hinge bricks were accomplished by manually moving the locations of staple crossovers to avoid large gaps in the connected helices without a crossover. Staple crossovers also had to be manually adjusted to avoid sandwich strands or kinetic traps which were not excluded by the autostaple feature.

RMSD values were plotted with each simulation step for H1 and H2 structures on MrDNA cryogenic electron microscopy (cryo-EM) reconstruction [13]. The RMSD plot indicates the average distance the residues of the structure have deviated from their original positions since the simulation began and was used to determine whether the simulation had converged. MrDNA displayed the structure at equilibrium once the simulation was complete, indicated by plateau in RMSD across simulation steps (Supplemental Fig. 3). For our structures, the default ten million step simulation was deemed sufficient for predicting the equilibrium structures of H1 and H2 to fold as expected, converging around the 5 millionth step. The simulation results displayed a hinge structure with two distinct brick components connected at the pivot point by single stranded scaffold regions, as designed in caDNAno (Fig. 4). The drift of the residues over time in the structure are also visible on the Visual Molecular Dynamics (VMD) interface, which displays the structure at regular intervals of simulation steps.

**Figure 4.**
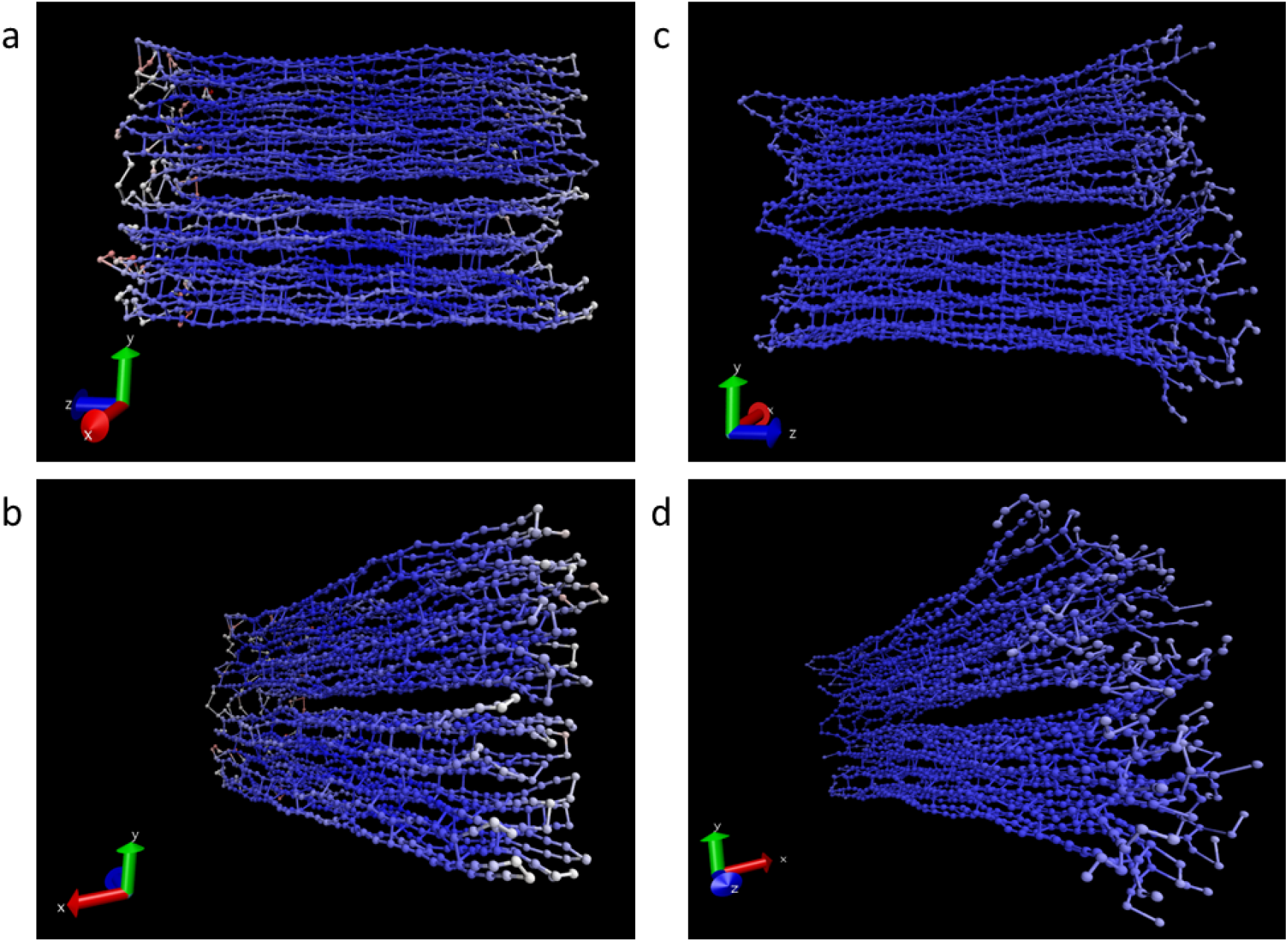
3D Simulations of H1 and H2. (a) Side view of H1 structure simulated on MrDNA visualized on Visual Molecular Dynamics (VMD) software. (b) Side view of H2 structure simulated on MrDNA visualized on VMD. Blue, white, and red represent high, intermediate, and low RMSF values, respectively.

MrDNA simulations displayed higher fluctuations at the free ends of H1 and H2. The increased fluctuations of residues at the ends of the hinge structure are expected. Residues at the ends of the hinge will move a greater distance relative to residues closer to the pivot point albeit the same angular movement. Both MrDNA and CanDo can output a range of RMSF values that were computed. The ranges were 0.91 - 4.30 nm and 0.62 - 4.53 nm computed by MrDNA. These provide a quantitative measure of how each residue within the nanostructure fluctuates, and are complementary to the heat maps provided by the simulation programs [39], [41], [42]. The close overlap of RMSF values and predicted structure from two different simulation programs increases the likelihood that the structures will form as designed.

### Hinge Assembly

Previous studies using agarose gel electrophoresis to determine DNA nanostructure formation have shown that a shift in band migration relative to pure scaffold are indicative of structure formation, yet nanostructure migration relative to scaffold controls can be dependent on the structure itself or the assembly conditions [6], [11], [43]. To determine whether staples were bound to the scaffolds of H1 and H2 as expected, gel electrophoresis was used to resolve differences in gel migration patterns between scaffold and hinge assemblies. An upward shift (i.e., slower migration) in the band location of H1 and H2 suggested staples strands were indeed being bound to their respective scaffolds, a requirement for proper structure assembly (Fig 4). A dark smear below 100 bp in the H1 and H2 lanes corresponded to excess staple strands, of which the majority could be removed with 2 washes during purification of H1 and H2 (Supplementary Fig. 7).

In addition to gel electrophoresis, transmission electron microscopy (TEM) was performed on purified H1 and H2 hinges to verify whether the structures assembled into the intended hinge shape. Images revealed hinge-shaped structures depicting two DNA bricks connected at one end and opening of varying angles from the connection point. Measurements of hinge lengths and angles from TEM images gave a mean brick length of 38.2nm ± 8nm and a mean angle of 96.2 degrees ± 20.7 degrees. Brick measurements on average differed by 11.98 nm compared to our predicted length of 50 nm.

H1 and H2 hinges were then incubated together to evaluate their ability to polymerize into HONs. Bands at the same location as H1 and H2 single hinges (Fig. 5b) persisted after HON assembly, indicating that some single hinges remained and had not polymerized. A minimum of five additional bands could also be visually detected, suggesting at least five different nanostructures had formed (data not shown). These were suspected to represent a stepwise increase by one hinge to the complex (e.g., H1–H2 for the second lowest band, H1–H2–H1 for the third lowest band, etc.). As the order of the species increased, band intensity decreased. This suggests a decreasing probability of formation of longer stable nanostructures as expected by thermodynamic limits [44]. Any higher order structure formation beyond six hinges (represented by the highest band) may not have been detected due to the concentration falling below the detection limit on the gel.

**Figure 5.**
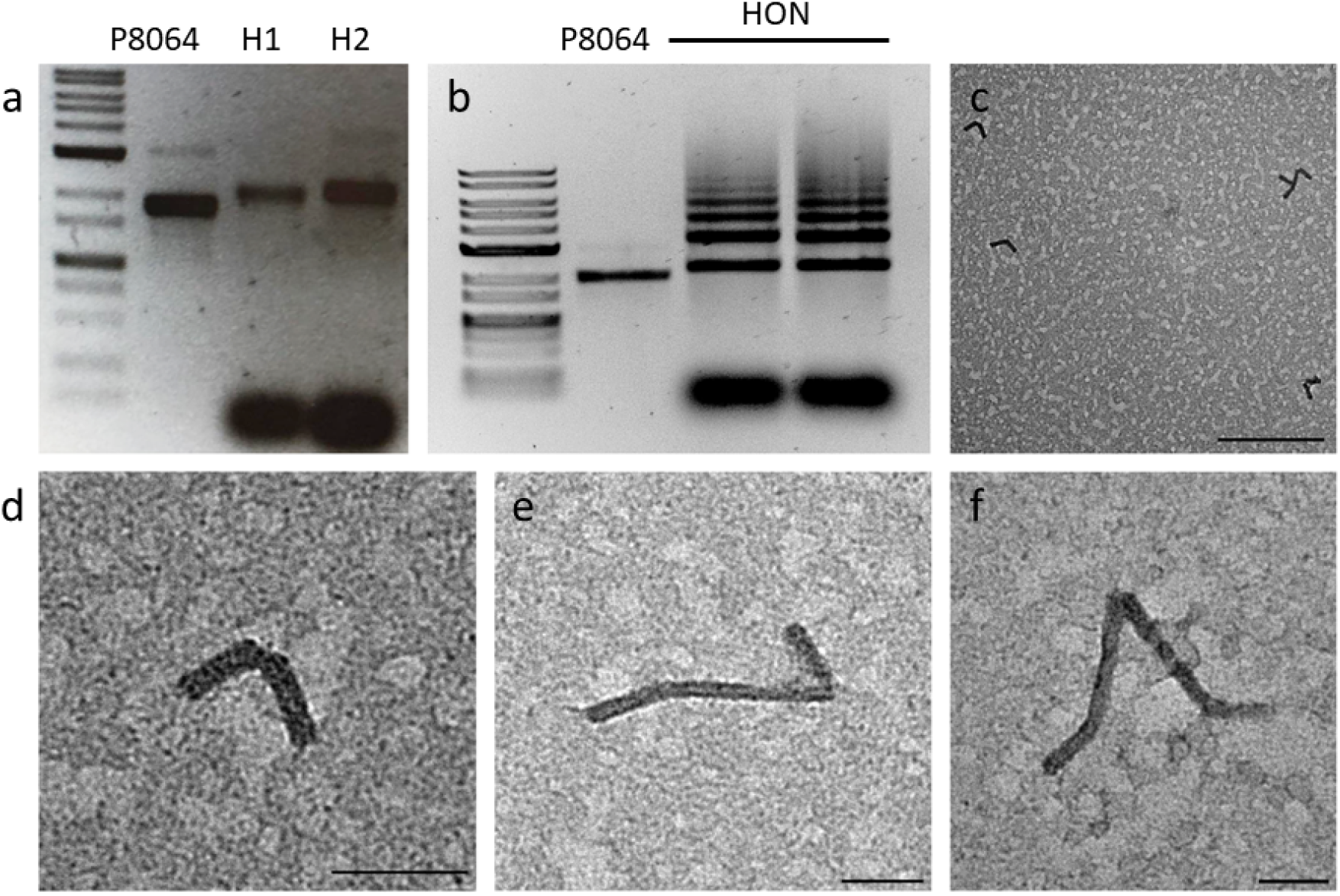
Formation Singe Hinges and Higher Order Hinges. (a) H1 and H2 post assembly. Hinges were formed with a 5:1 staple to scaffold molar ratio in 20 mM MgCl_2_ with a 37-hour annealing ramp. Gel order: 1 kb DNA ladder, p8064, H1, H2 on a 1% agarose gel. (b) HONs post assembly. HONs were formed with purified H1 and H2 incubated in the presence of polymerization and neighbour strands in 20 mM MgCl_2_ with a 21-hour annealing ramp. Gel order: 1 kb DNA ladder, p8064, two independent HON assemblies on a 1% agarose gel. (c) H1 TEM image. Scale bar is 200 nm. (d) H1 TEM image. (e) Dual-hinge TEM image. (f) Triple-hinge TEM image. TEM images are representative of three sample preparations and multiple images obtained. Scale bars are 50 nm.

TEM was used to examine the formation of HONs. The number of combined hinges within a HON was determined from the known dimensions of H1 and H2. The end of each hinge within the EHS formed a straight path, demonstrating structurally rigid end-to-end connections. The average total length of HONs visualized was 147.36nm ±15.77 nm, with the dual hinge complex yielding 96.41nm and the triple hinge complex yielding a length of 223.02 nm (Figure KB). The polymerized lengths of the HONs were determined to be discrete multiples of the length of a single hinge structure.

### Locking Mechanism

To determine whether each strand of the locking mechanism could first recognize its target strand in the sequence, strands and their targets were incubated together. Resulting hybrids were identified using SDS-PAGE based on shifts in band location compared to negative control lanes 1-5, representing singular components of the locking mechanism (Fig. 6). One band between 50 and 75 bp was present in the lane containing the padlock and strand. Its alignment with the padlock control suggested that all locking strands were bound to the padlock. Due to the fainter appearance of the locking strand, a gradient of lock to padlock strands were incubated to further confirm the lock strand could bind to the padlock (Supplementary Fig. 8). When padlock, lock, and antilock were incubated together a band was produced between 25 and 35 bp, which demonstrated the lock and antilock strands hybridized together. The appearance of this hybrid in the presence of unbound padlock suggested that the lock and antilock strand had a higher affinity as expected and that the padlock did not interfere with binding. A similar trend was observed in the incubations of the key strand and the padlock strand. In the presence of the lock strand, a key and padlock hybrid were observed between 75 and 100 bp. Persistence of a band between 50 and 75 bp indicated that not all padlock strands hybridized with key strands despite their sequences having a greater degree of complementary compared to either lock strand sequence. This could be due to an equilibrium between bound and unbound key-padlock strands or a slightly nonoptimal molar ratio. The presence of a band between 35 and 50 bp indicates the presence of key binding to antikey. Finally, when lock and key strands were incubated together in the presence of their counterparts the persistence of their respective hybrids demonstrated that the combination of all locking mechanism strands did not interfere with each strand’s ability to recognize and bind to its counterpart.

**Figure 6.**
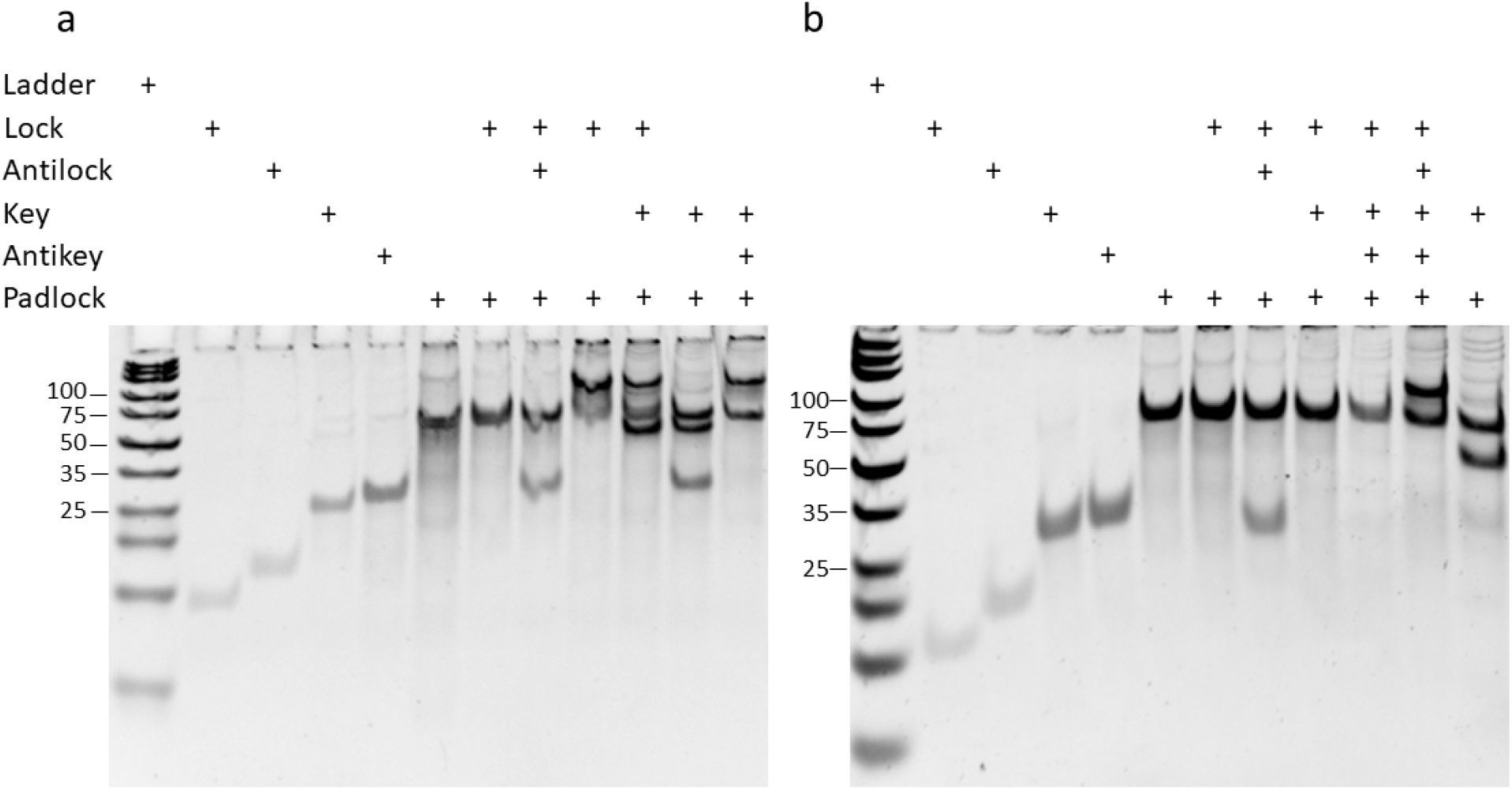
Locking Mechanism strand recognition and displacement. (a) 20% Polyacrylamide gels demonstrating locking mechanism strand recognition for H1 padlock 2. Shaded cells in the table represent the strand components incubated in each condition. All component strands for each gel are the same aside from the padlock strand as indicated. (b) Demonstration of the ability of the locking mechanism strands to displace their intended targets through the addition of either the antilock, key, or antikey strand for H2.

After determining capacity of the locking mechanism strands to recognize their counterparts, capacity to displace their intended strand in the mechanism sequence was also evaluated using the same approach. The band pattern suggests that after incubation with the complement strand, the antilock, lock, and key were able to displace their target from the padlock strand as indicated by the presence of their respective hybrids following the second incubation with padlock hybrids and either the antilock, key, or antikey strand (Fig. 6b). Consistent with the results in Fig. 6a, the lock and antilock hybrid was observed between 25 and 35 bp, the key and padlock hybrid between 75 and 100 bp, and the key and antikey hybrid between 35 and 50 bp. The appearance of additional bands such as excess padlock or those found at non-hybrid or control locations suggest a need for further locking mechanisms sequence optimization as discussed below. The same pattern was observed in the gels for both padlock strands of H1 and H2 (Supplementary Fig. 9).

Following assessment of the locking mechanism in isolation, its effectiveness in inducing conformation changes in H1 and H2 was investigated. In each sample condition (hinge, hinge + lock strand, hinge + key strand), a range of angles from 0 - 180 degrees between the hinge bricks were observed under TEM imaging. The number of hinges in either a closed conformation (0 degrees) or open (> 0 degrees) was summed to determine changes in the ratio of closed to open hinges across the different conditions. Compared to the control, hinges incubated with the lock strand exhibited the highest proportion of closed hinges, followed by the control and incubation with the key strand. Conversely, hinges incubated with the key strand had a significantly lower proportion of closed hinges compared to both the control and lock incubated hinges (Fig. 7). This matched expectations, as the lock strand was designed to lock the hinges in a closed conformation, and the key strand, an open conformation. Skewing in distributions of the key and lock conditions toward their intended conformations was also observed. The magnitude of the median ratio values less than 0.5 indicated that greater optimization of the locking mechanism is warranted to ensure the majority conformation of the hinge per conditions results as intended.

**Figure 7.**
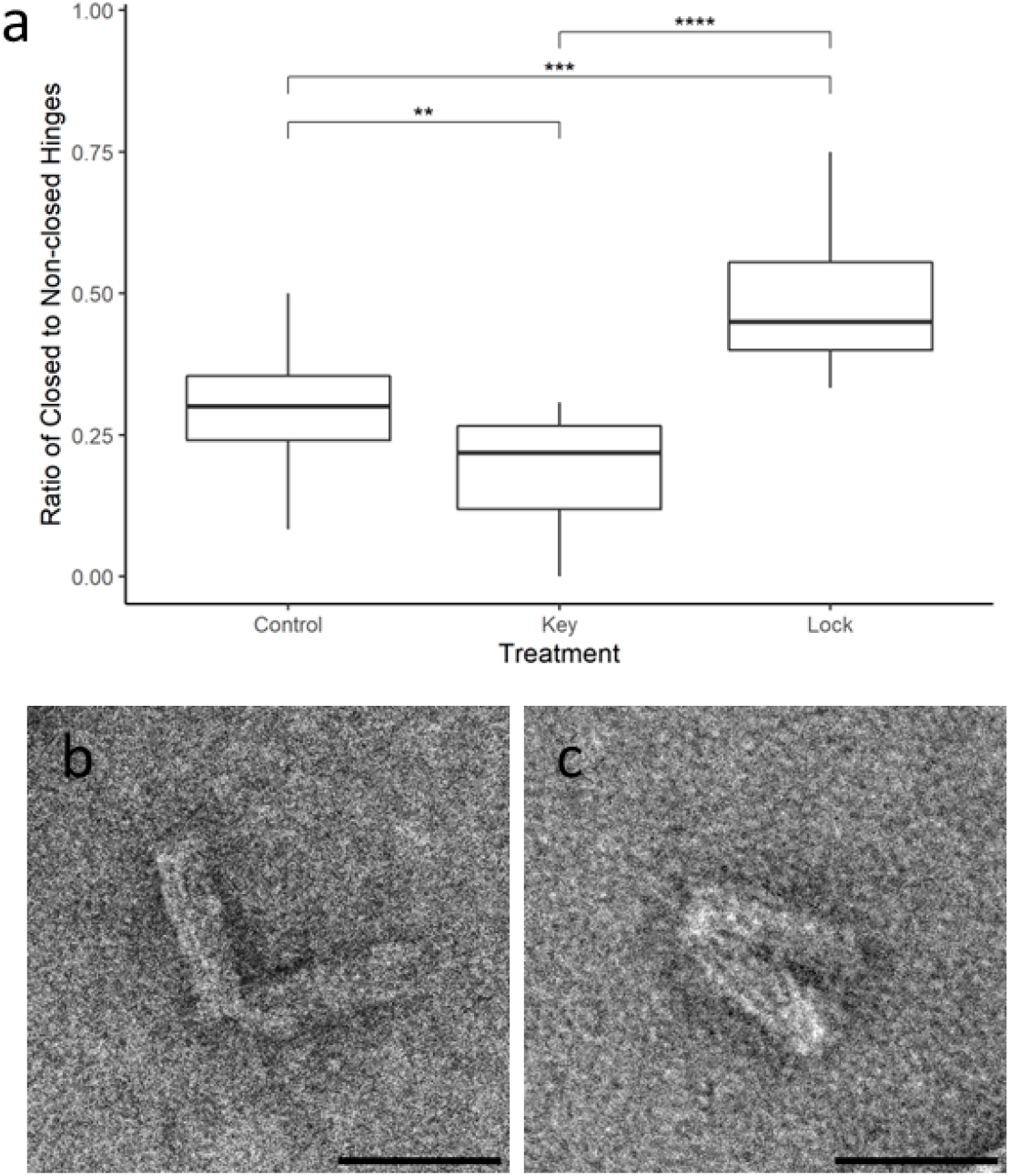
Incubation with lock results in a higher proportion of closed hinges. (a) Distribution of closed to non-closed hinges when treated with key strand, and lock strand, or no treatment (**P< 0.01, ***P<0.001, ****P<0.0001). Data was analyzed with a one-way ANOVA followed by Tukey multiple pairwise comparisons test. Hinges in the fully extended state were not included in determining the proportion of closed hinges for each condition. (b) TEM of closed hinge. (c) TEM of open hinge. Scale bars are 50 nm.

## DISCUSSION

This work introduces a self-assembling DNA nanohinge system capable of reversible conformational changes, and higher-order structure formation. Assembly of these more complex nanostructures was achieved using two hinge-shaped nanostructures capable of binding brick to brick in an alternating series to scale the length of the overall structure. The incorporation of additional ssDNA sequences employing TMSD served as the mechanism for inducing open and closed states in single hinge units.

When using the scaffold-based approach of DNA origami to assemble DNA nanostructures, staple strands typically occupy a specific location on the scaffold, corresponding to a unique staple sequence, that causes the scaffold to bend into the desired shape [9]. However, as the complexity and size of structures increase, so do the number of staple strands required, which can increase staple sequence overlap and prevent efficient binding of staples to their intended scaffold location. While H1 and H2 increased the number of basic units needed to assemble higher order nanostructures, two different hinges were created to simplify the process of creating hinge arms that allow an interlocking fit. This was necessary to avoid crossovers around the blunt-end region that could deform end-end conformation, increasing the risk of structure aggregation [45]. Further, this approach was taken to conserve as many natural crossover events in the scaffold strand as recommended by caDNAno. In doing so, each interconnecting helix had to be manually checked to ensure that the directionality of polymerization strands was maintained and could bind the ends of two bricks together.

The lack of nanostructures in earlier H1 and H2 iterations, prompted several design changes that led to the success of the current H1 and H2 design. First, decreasing the number of helices in each brick from 36 to 28 decreased their complexity in the number of interactions required between hinge bricks and in the number of staples required. In doing so, this decreased potential staple strand overlap and blunt end occurrences, as mentioned above. Furthermore, rather than using staple strands to connect two separate DNA bricks together at one end to form the pivot point, the current design utilized a single scaffold strand that ran through the hinge bricks and pivot point. It was suspected that these changes were necessary to minimize the risk that hinge bricks could not connect at the pivot if an excess of staple strands outcompeted the binding of specialized staples for this connection. However, previous studies have reported the successful assembly of hinge-shaped nanostructures using this approach [5], [6]. Furthermore, the use of a single scaffold strand in its entirety, minimized unbound sections of the scaffold which may interfere with the specific binding of staple strands during structure assembly. Further iterations of H1 and H2 could be made by decreasing the number of helices, thereby further decreasing the number of staples required for structure assembly. A decreased cross-sectional area may increase structure flexibility and decrease the number of connection points between H1 and H2 [46]. The changes undertaken to produce the current iterations of H1 and H2 predicted stable structures at the simulation stage which were verified at the assembly stage.

There are several considerations regarding the reproducibility of the DNA nanostructure design process described. First, there is an element of subjectivity and variability in the design process using caDNAno. Although caDNAno provides a useful framework in which to design DNA nanostructures, critical details may vary depending on the user. caDNAno allows users to either manually decide where to place staple strands in relation to the scaffold strand or to use an autostaple option. However, the autostaple option does not detect and remove the occurrences of kinetic traps, and therefore must be manually resolved. Whether this variability has any impact on the stability or likeliness of the structure to form is unknown, and thus presents a limitation to our design pipeline. This makes any quality control or validation of the 2D design difficult, as there are no standardized benchmarks for comparison. For instance, in our staple strand sequence similarity analysis, we saw that staple strands had an average of 9.8 sequential base regions that were identical to another staple strand. Although there are a variety of possible algorithms to compare the similarity of two sequences, we chose the number of sequential nucleotide bases that were found in two or more separate staple strands because it is a simple metric that considers both the entire staple sequence and the sequence of the separate binding domains on the scaffold strand.

The interaction between the number of identical base regions and potential for competitive scaffold binding or increasing the likelihood of strand aggregation is uncertain. A detailed study of the thermodynamics of the DNA nanostructure formation process may provide such insights, and how sandwich strands or kinetic traps affect the probability of proper structure formation. As a result, the structure validation was done using 3D simulation software. An important consideration that arises from relying on 3D simulation software to validate the designed nanostructures is the reliability of the simulation software. The primary 3D simulation method used for our design pipeline was designed to shorten the time to validate the formed DNA nanostructures from the traditional method where nanostructure validation would have been done iteratively by verifying lab results. This simulation framework was verified by comparing cryo-EM reconstruction images and designed nanostructure shapes. For the purposes of our study, this approach was deemed sufficient to validate our nanostructure designs.

Higher order structure assembly of multiple H1 and H2 units was observed, further validating our design process. Although TEM showed that our conditions led to successful assembly, a gradient of agarose gel band intensities suggested that shorter HONs containing < 3 hinges were more likely to form than longer ones up to 6 hinges in length. Consistent with this observation, HONs of two to three hinges were observed more frequently in TEM images than higher-order ones. Beyond this we could not resolve bands within the limit of detection of our gels [47]. Increasing incubation time may increase the length of HONs by prolonging the interaction time between H1 and H2 [10], [11]. As such, incubation time is expected to be a contributing factor in controlling HON length and increasing the concentration of longer HON structures. Additional steps to detect the presence of HONs > 6 hinges may be to run lower percentage gels (< 1%) for a longer duration of time [48]. Iinuma et al. observed a similar intensity gradient for higher order polyhedra formed by the polymerization of tripods, which were run on a 0.8% agarose gel [23]. Alternatively new dyes have been developed with improved sensitivity for DNA detection [47]. Size exclusion chromatography may also prove a more effective means of characterizing different HON structures formed and has been previously used in the characterization of other nanostructures [49].

Assessment of the locking mechanism, both isolated and incorporated with single hinges, suggested that strands of this mechanism were able to recognize and displace their intended targets, as well as induce conformational changes in the DNA nanohinge. The persistence of excess padlock strands in conditions in which greater padlock hybrids were possible, such as key and padlock hybrids, suggests that a 1:1 molar ratio may not be sufficient. This trend could extend to other strands of the locking mechanisms and their counterparts. For instance, key strands may not be able to displace all of the lock strands that have been bound to padlock strands as intended. A higher ratio or greater degree of complementarity between locking mechanism strands may be required to minimize the presence of unhybridized strands that would decrease the efficiency of the locking mechanism. Despite consistent loading, bands representing lock and antilock strands appeared the faintest. In conditions where unbound lock is expected, the amount of unbound lock strand, not including those that did not hybridize, may be below gel detection limits [47]. Moreover, the appearance of additional bands in some conditions not explained by the appearance of hybrids suggests that there may be undesirable sequence overlap leading to off-target binding of locking mechanism strands. This consideration does not include the sequence overlap found in overhangs to initiate TMSD. Future approaches include redesigning portions of the locking mechanism strands to improve binding affinity such as by increasing sequence length and scanning for potential off-target binding. Additionally, incubation conditions, such as temperature, could be optimized to favor annealing of the next strand in the locking mechanism sequence.

Incubation of hinges with the lock strand and the key strand demonstrated the highest and lowest ratio of closed to open hinges respectively. This suggests that the locking mechanism was able to induce conformational changes in single hinge units. To simplify interpretation it was assumed that all hinges that had an angle of 0 degrees in TEM images were closed hinges as there were no distinguishable features between hinges closed due to the locking mechanism and those deposited onto TEM grids in that conformation by chance. It was hypothesized that if the locking mechanism was able to induce conformational changes that this would be reflected in the proportion of closed hinges across the different incubation conditions, as was observed. Although closed hinge proportion trends matched expectations, the median ratio of closed to open hinges in hinges incubated with the lock strand was less than 0.5, suggesting a low efficiency in inducing the hinges to adopt a closed conformation.

There are many variables with potential impact on assembly and function of the locking mechanism. As such, verifying incorporation of the padlock strands in H1 and H2 was nontrivial because excess staple strands that persisted post purification obscured the expected band location of the padlock strands. The most influential part of the design is the assigned location of the padlock strands due to their impact on the ability of the hinges to open and close. A balance between sequence length and optimal tension of the strands needs to be established. For example, placing the strands closer to the conjunction would require shorter sequences. However, they would be subjected to higher stress to transition between conformations. In contrast, excessive elongation of the padlock strands hinders their binding efficiency during hybridization due to the complex binding kinetics such as increased entropy and non-specific binding [50], [51]. Finally, the possible presence of “leaky” systems, where there is an absence of an input strand, can also disrupt the behavior of strand displacement cascades [52].

## CONCLUSION

Recent advancements in DNA nanotechnology have paved the way for our nanohinge system to be used in biosensing [53], [54] and nanofabrication [21], [55], [56]. For applications in biosensing, the detection of a target nucleic acid through displacement of the lock strand could change a hinge from a closed to open conformation, rendering it ideal for fluorescence-based detection systems. This could enable the development of highly sensitive and selective biosensors capable of detecting multiple targets simultaneously through an array of hinges sensitive to different targets. Higher order nanostructures could extend the targeting scope to simultaneously detect other biomolecular targets, triggered conformational changes such as the opening of a DNA nanobox [57] and the activation of a nanorobot [54] upon protein binding. For nanorobotics, single hinges or HONSs could be used as dynamic joints to aid in structure movement [6] or molecule localization [55], [56]. Collectively, nanohinges can serve as pivotal constituents in nanorobotic construction by facilitating controlled motion and fine manipulation of molecular components.

The scalability of self-assembling DNA nanohinges makes them amenable to micrometer-scale structure fabrication. By using single hinges as parts in bottom-up fabrication of more complex structure designs, the risk of kinetic traps and misfolded structures will be minimized. Future work could integrate alternative actuation methods, including paramagnetic beads [6], light-responsive [32], [58], or enzymatic-based approaches [56], [59] to expand the functional versatility of our DNA nanohinge system.

## ACKNOWLEDGEMENTS

This work was performed under the auspices of the Natural Sciences and Engineering Research Council (NSERC) of Canada and the Canada Foundation for Innovation (CFI). ML, CS, FI, and SN were supported by the NSERC CREATE Ecosystem Services, Commercialization Platforms and Entrepreneurship (ECOSCOPE) training program at the University of British Columbia. Additional funding was provided by the Professional Activities Fund through the Dept. of Applied Sciences at the University of British Columbia. We would like to express our deepest appreciation to Dr. Stephanie Lauback, Dr. Dan Bizzotto, Dr. Alexander Marras, Adrian Jan Grzedowski, Resmi Radhamony, Jennifer Bonderoff, Derrick Horne, Marlene Chow, and all other members of the project team for their unwavering dedication, expertise, and insight, and to the undergraduate Biomolecular Design competition which served as human scaffold for this work.

## Notes

### Competing Interest Statement

The authors have declared no competing interest.

### Summary of Updates

Supplemental file updated.

