## Supplementary Materials for "Self-Assembly of a Repeatable DNA Nanohinge System Supporting Higher Order Structure Formation"

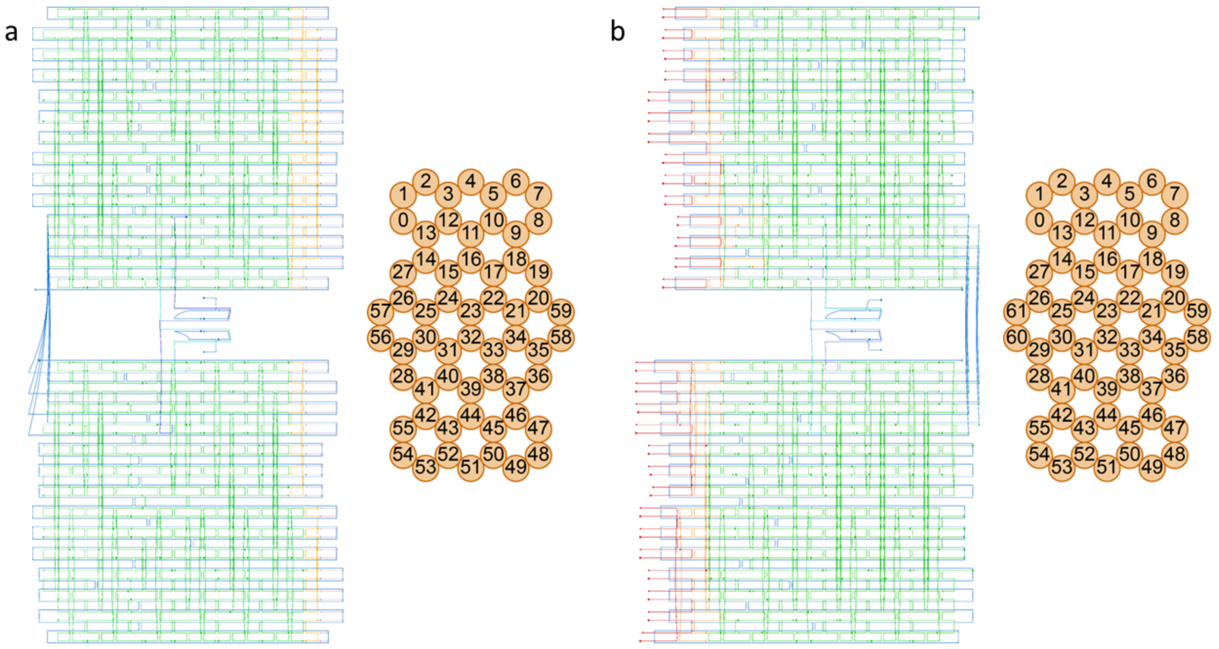


**Supplementary Figure 1.** CaDNAno design of hinges. (a) H1. (b) H2. P8064 scaffold is shown in blue, core staples are in green, and polymerization strands with overhangs in red. Padlock strands are shown in cyan and purple. The location of polymerization strand binding sites are shown in red. Cross-section of  H1 and H2 hinges show the locations of the double helix bundles constituting each structure.

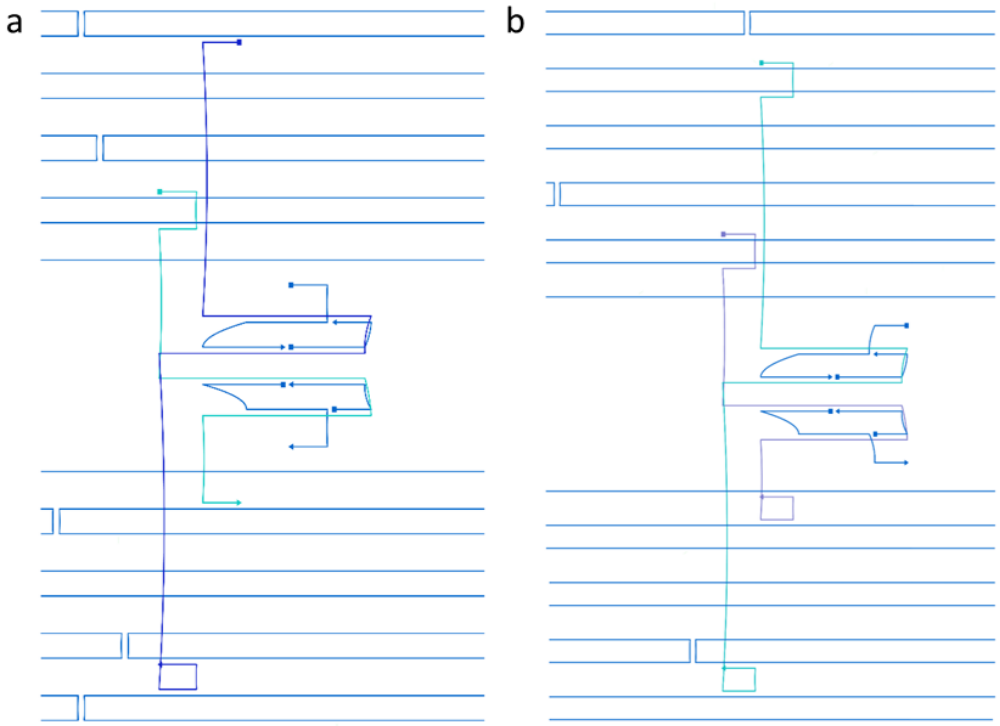


**Supplementary Figure 2.** CaDNAno design of locking mechanism. (a) H1. (b) H2. P8064 scaffold is shown in blue. Padlock strands are shown in cyan and purple.

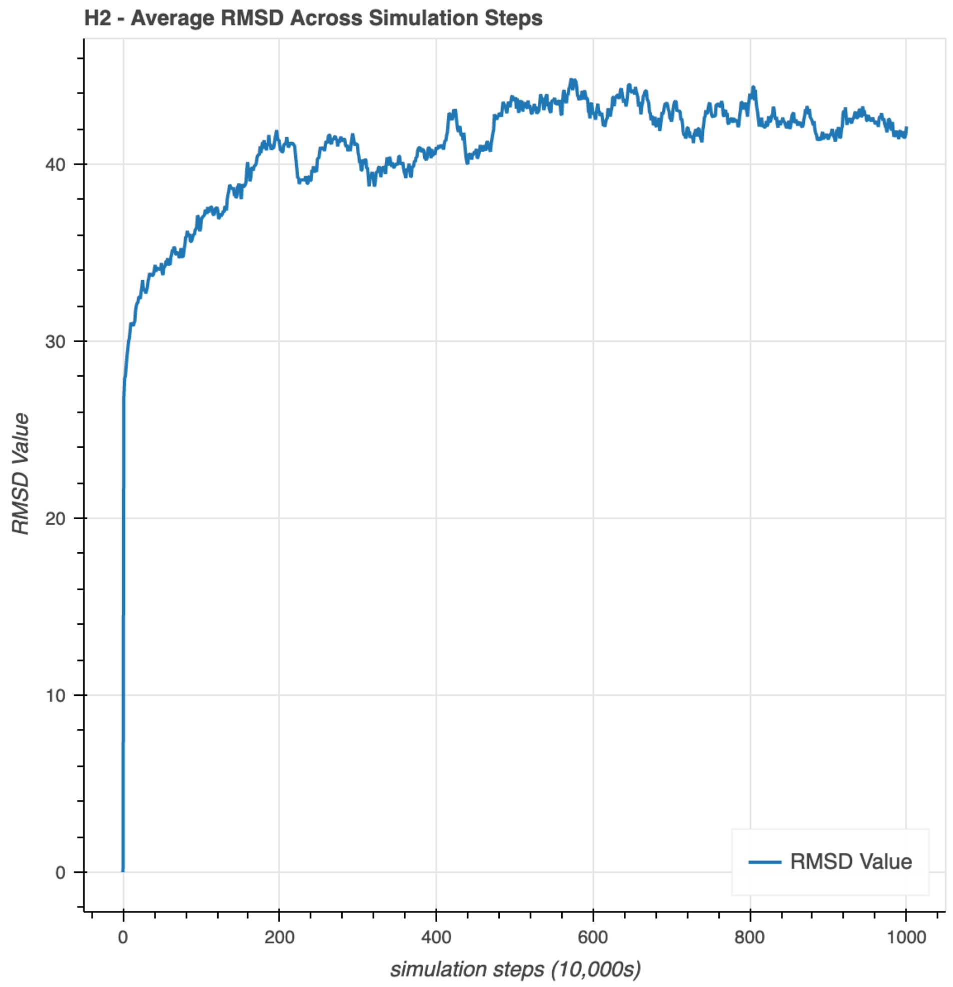


**Supplementary Figure 3.** The graph of the RMSD Values for H2. RMSD Values across simulation steps were obtained for 10,000,000s in simulation time. It was assumed that the prediction of the equilibrium H2 structure was complete when the average RMSD converged at 5,000,0000s, showing no significant fluctuations from 5,000,000s -10,000,0000s. The same outcome was obtained for H1.

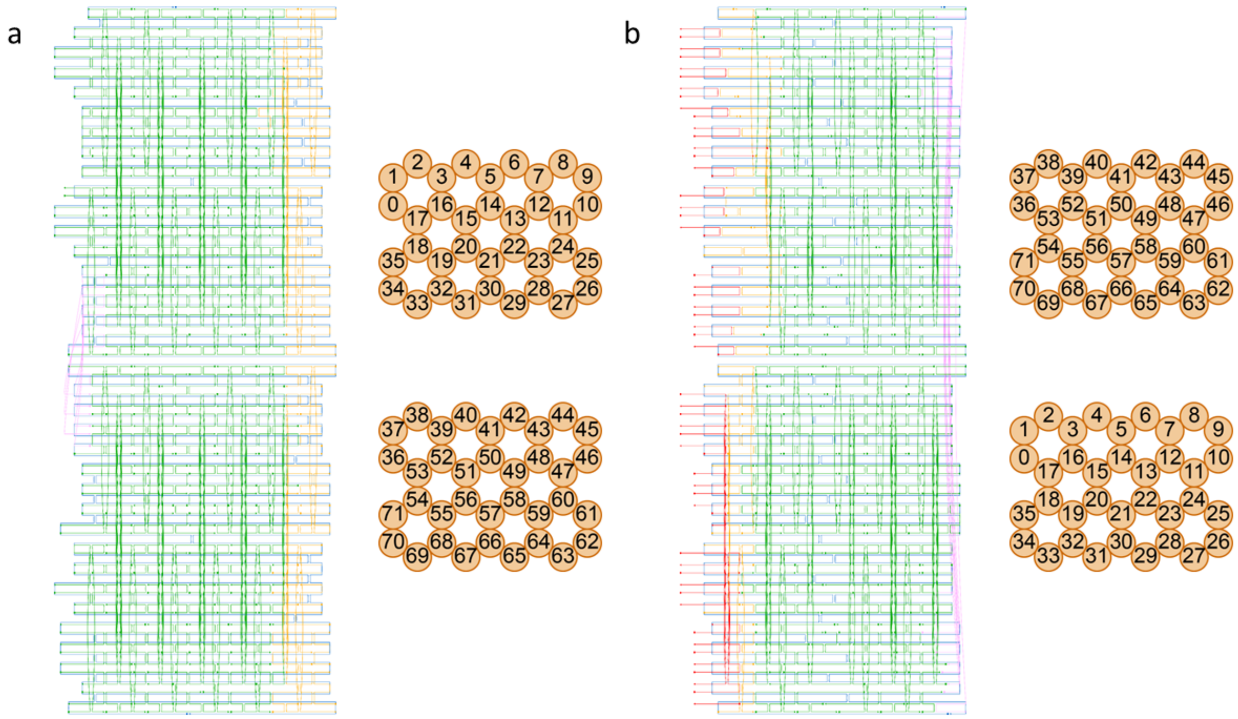


**Supplementary Figure 4.** Previous CaDNAano design and visualization of hinges and their cross-sections. (a) H1'. (b) H2'. P8064 Scaffold is shown in blue, core staples are in green, and polymerization strands with overhangs in red. Padlock strands are shown in light blue and purple. The location of polymerization strand binding sites are shown in red. Cross-section of H1' and H2' hinges show the locations of the double helix bundles constituting each structure.

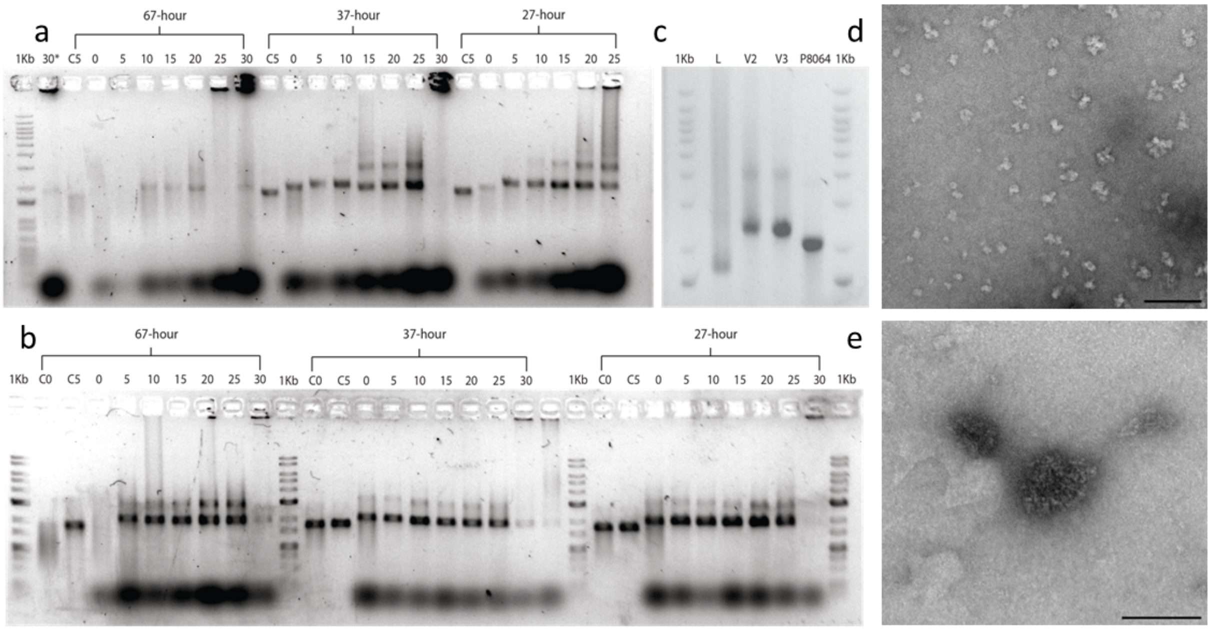


**Supplementary Figure 5. Assembly results of previous hinge designs. (a)** H1' and **(b)** H2' salt and annealing ramp screens. Hinges were formed at a 5:1 staple to scaffold molar ratio in 0 – 30 mM $MgCl_2$ with a 27, 37, and 67-hour annealing ramp and run on a 1% agarose gel. The H1' 30 mM $MgCl_2$ sample for the 27-hour annealing ramp is indicated by 30*. C0 and C5 denote p8064 scaffold controls supplemented with 0 and 5 mM $MgCl_2$, respectively. **(c)** H1' and H2' assembly compared to Lauback et al.'s hinge (L). H1' and H2' were formed in 15 mM $MgCl_2$ with a 37-hour annealing ramp and run on a 1% agarose gel. The 1 Kb DNA ladder used in (a) and (c) was from Thermo Fisher, whereas in (b) was from FroggaBio. (d) TEM imaging of H1'. (e) TEM imaging of H2'. No hinge-like structures were observed in TEM images of H1' and H2'.

Images were representative of three sample preparations and multiple images obtained. Scale bars are 100 nm. Images obtained from Law, M, 2021. *From Nano to Micron Scales: The Construction of a Scalable and Self-polymerizing DNA Origami Hinge.* Thesis. (BSC). University of British Columbia .

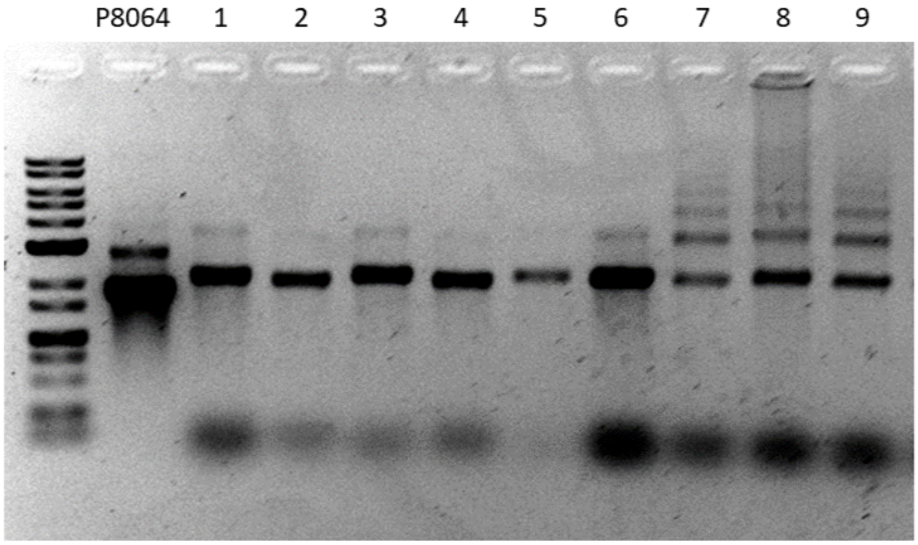


**Supplementary Figure 6.** Hinge Assembly Parameter Optimization. H1, H2, and HON assembly were prepared under different assembly conditions that varied the concentration of $MgCl_2$ and TE vs. TAE buffer. Order of 1% agarose gel: 1 Kb DNA ladder, P8064 scaffold, 1. H1 in 10mM $MgCl_2$ TE, 2. H1in 15 mM $MgCl_2$ TE, 3. H2 in 15 mM $MgCl_2$ TAE, 4. H1 in 15 mM $MgCl_2$ TAE, 5. H2 in 20 mM $MgCl_2$ TAE, 6. H2 in 20 mM $MgCl_2$ TE, 7. HON in 20 mM $MgCl_2$, 8. HON in 10 mM $MgCl_2$, 9. HON in 15 mM $MgCl_2$. HON conditions were assembled in TE buffer. Assembly in 20 mM $MgCl_2$ provided less smearing during HON assembly.

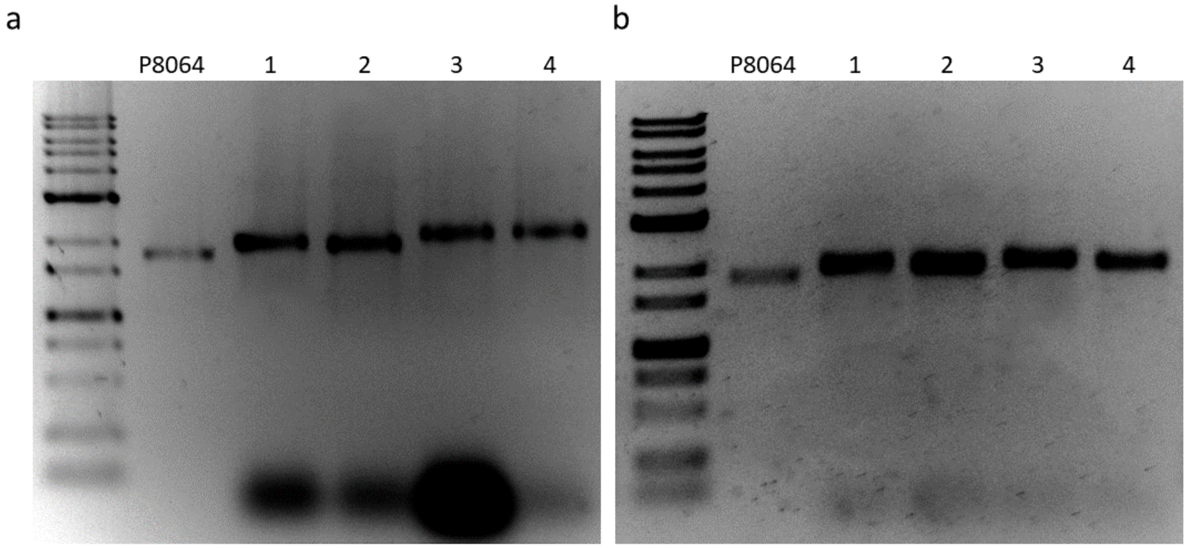


**Supplementary Figure 7.** Purification of nanostructures. (a) H1 and H2 post incubation. (b) H1 and H2 post purification using 100 kDa Amicon Ultra Centrifugation filters. Hinges were assembled with 1:2 or 1:5 molar ratios of padlock 1 and padlock 2 strands to scaffold.
Order for both 1% agarose gels: 1 Kb DNA ladder, P8064 scaffold, 1. H1 at 1:2 scaffold to staple molar ratio, 2. H1 1:5, 3. H2 1:2, and 4. H3 1:5. Excess staple strands located below 100 bp were notable absent post purification.

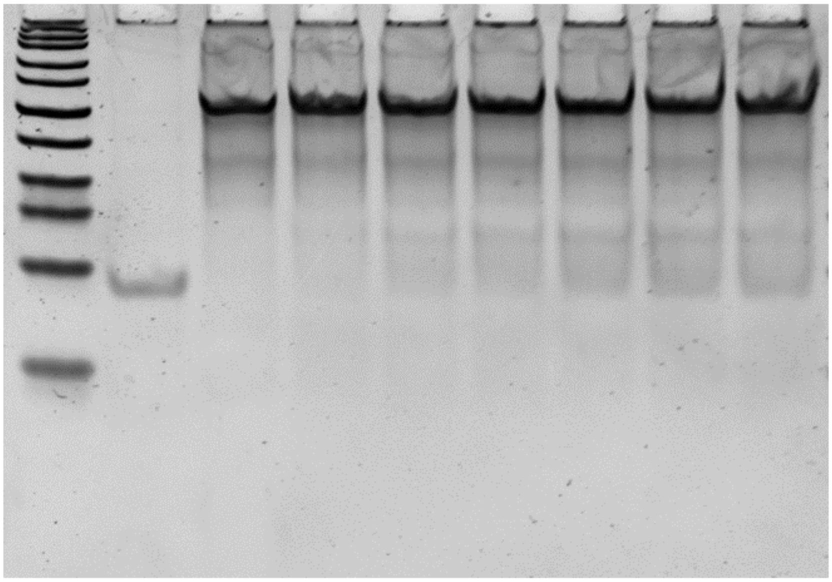


**Supplementary Figure 8.** Binding of lockstrand to padlock strand. To that confirm lock strands could bind to padlock strands, increasing molar ratios of lock strand was incubated with H2 padlock 1 from 1:0.5 to 1:3 and incubated at 30°C for 25 minutes. Excess lock strand was observed at higher molar ratios suggesting saturation of complementary binding sites on the padlock strand. Due to the same sequences of all padlock strands aside from their overhangs, the same result was expected of all padlock strands.

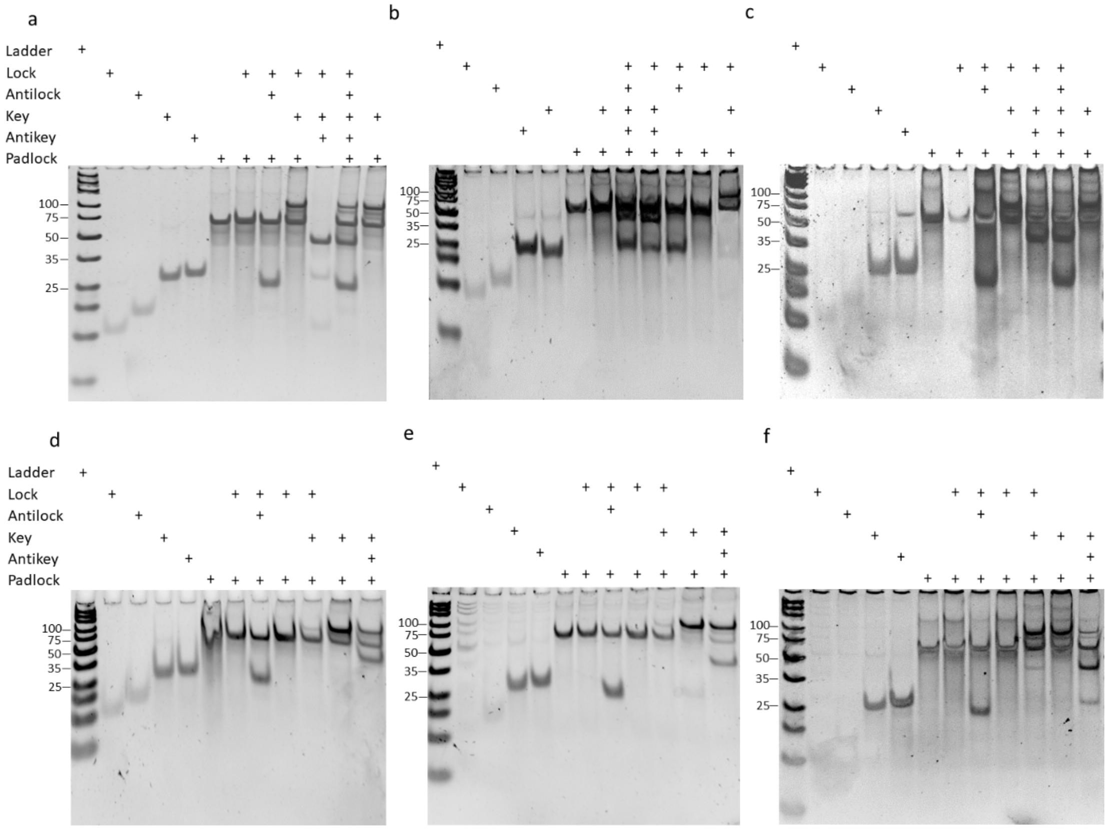


**Supplementary Figure 9.** Panels a-c represent 20% Polyacrylamide gels demonstrating locking mechanism strand recognition for H1 and H2 hinges. + represents the presence of strands incubated in each condition. All components strands for each gel are the same aside from the padlock strand as indicated. (a) H1 padlock 1 (b) H2 padlock 1 c) H2 padlock 2. Panels d-f demonstrate the ability of the locking mechanism strands to displace their intended targets through the addition of either the antilock, key, or antikey strand. (d) H1 padlock 1. (e) H2 padlock 1. (f) H2 padlock 2.

### Supplementary Tables

**Table S1 List of ssDNA Staple Classifications, Definitions and Colors for the nanohinge.** This table provides an overview of the different classes of single-stranded DNA (ssDNA) staples used in the nanohinge, along with their definitions and associated colors. The locking mechanism strands and anti-locking strands, are used to bind to PADLOCK1 and PADLOCK2 and do not have specific colors. The color legend serves as a reference for Supplementary Figures 1 and 2, which show the appearance of the different staple classes.

| ID | Color | Definition |
|---|---|---|
| CORE | green | Core Strands serve in maintaining the structural integrity of the H1/H2 scaffold. |
| NEIGH | orange | Neighbor strands facilitate the formation and maintenance of a H1 and H2 hinge-pair structure. |
| POLY | red | Polymerization strands have overhang sequences that can join a H1 and H2 hinge-pair with another pair. (note: located on H2 only) |
| PADLOCK1 | Purple | The two padlock strands that are bound to each side of the hinge in a H1 and H2 pair. |
| PADLOCK2 | cyan | |
| LOCK | | The locking strand binds to the padlock and the strand that force the H1 and H2 to adopt closed conformation. |
| ANTI-LOCK | | The anti-lock strand displaces the lock staple strand during the opening of a H1 and H2 hinge. |
| KEY | | The key strand is the sequence that binds to the padlock strand, allowing a H1 and H2 hinge to prop open. |
| ANTI-KEY | | The anti-lock strand displaces the key strand during the closing of a H1 and H2 hinge. |

The following sequences were used in the formation of the H1 and H2 hinges. The Start and End columns denote the helix[base] positions at the 5' and 3' ends of the DNA staple strands in the CaDNAno design for their respective designs.

**Table S2A. List of Core and Neighbour Strands (H1)**

This table denotes the ssDNA core and neighbour staples sequences used for the H1 hinge. The Start and End columns denote the helix[base] positions at the 5' and 3' ends of the DNA staple strands in the CaDNAno design for the respective designs (Supplementary Fig. 1A). Refer to Table 1 for the definitions of the IDs.

| Start | End | Sequence | Bases | ID |
|---|---|---|---|---|
| 1[70] | 23[76] | AAACCGAAAGTTTAGGGGTTTCAGGC GGTCTGTAAATCGTCTGTCACAC | 49 | CORE |
| 28[125] | 40[112] | AAATCCCCCCAGCACATTAATGAATC GGACATTAAGGTAACG | 42 | CORE |
| 47[126] | 55[132] | AACAAGAGTCAATCTCGCATTAATAG GATGTAGATAGGGGACGAAAACG | 49 | CORE |
| 9[112] | 20[112] | AACAATATTTAACGTAGAACCTATCAT T | 28 | CORE |
| 1[91] | 23[97] | AACGGAAAACGCAATAACTATAACCT CCTTAATTTATACCTAGGGACAT | 49 | CORE |
| 8[90] | 12[77] | AACGGGTCAGCTAAAGAGAATGTATC ATGTGTGATAAATAAG | 42 | CORE |
| 45[49] | 47[62] | AACGGTGAGCAAAAATCGGTTGTACC AATCTAGCTTGCCGGA | 42 | CORE |
| 39[49] | 37[69] | AACGGTGGAGCCGCTCATTTGCCATA TTCCGATGCCAGCAGTTGGGCGG | 49 | CORE |
| 19[21] | 27[27] | AACTTTCTCCAGACACAACGCGGAAC CCACCGCCAAGGTGTATGAACCA | 49 | CORE |
| 54[83] | 32[77] | AAGATTAATCGTCATTCCGGCGCAAC TGAAACGACGTAAAGCCAGTGTC | 49 | CORE |
| 51[49] | 46[56] | AAGATTCATTCCATAAGAATTTACAGA CGTAATCT | 35 | CORE |
| 28[104] | 31[97] | AATAGCCAAGCGGTGCGGGGAGAGG CGGGCCTAAT | 35 | CORE |

| Start | End | Sequence | Bases | ID |
|---|---|---|---|---|
| 48[90] | 54[84] | AATCACCCTTTTGCACATCCATGCGAACTTTTAAAAACCAGACATCAAA | 49 | CORE |
| 37[91] | 48[91] | AATCCCGGCCCCAAAAGGCTAACAGTCA | 28 | CORE |
| 9[28] | 20[28] | AATCCTCTCGTCTTAACAGTTAGGAATT | 28 | CORE |
| 24[118] | 22[119] | ACAGGAAATAGAACCCTTCTGGTCTTTA | 28 | CORE |
| 17[77] | 7[83] | ACCAAGTTTACATCACATGTTATTAAACGTTTTTA | 35 | CORE |
| 30[55] | 28[42] | ACCAGTGAGTAAAGCCGATACGGGCAACACTATTAAAGAACG | 42 | CORE |
| 6[62] | 8[49] | ACCCACTTTTCATACCGCCACACCAGAG | 28 | CORE |
| 54[125] | 41[125] | ACCCTGACAGTTCAGACGACATACGCCA | 28 | CORE |
| 10[76] | 19[76] | ACCGACAATAAACAGGGAGAAAATCCTG | 28 | CORE |
| 39[77] | 35[83] | ACCTCACGGATCAAACTTAAATTGTGTACATCGACCAGCCTCGTGGTGC | 49 | CORE |
| 44[41] | 36[28] | ACCTGCTGGAACGATGAGGCTAGCATCACAATATCTGGTCAGAAAAGGATCGAGGT | 56 | CORE |
| 47[21] | 44[21] | ACGAGAAATCAACGACTGACCACTTAGC | 28 | CORE |
| 44[125] | 36[112] | ACGCCATACCAGTCGTGAAGGTTACACTCAGCATCAGCGGGGGCCAACGGGCAAAC | 56 | CORE |

| Start | End | Sequence | Bases | ID |
|---|---|---|---|---|
| 32[76] | 34[77] | ACTGCGCCCTGCGGCTGGTAAGCCGGGC | 28 | CORE |
| 54[62] | 52[49] | AGACTTCAATCGTTAACCCATATCGCGTCTGGAAGTTTCATC | 42 | CORE |
| 54[104] | 40[98] | AGCAAAGATCCCCCAAGATCGCGGTGCGTTTCCCA | 35 | CORE |
| 27[91] | 15[104] | AGCCCCCGGTCACGTTGATTAGTAATAACATGGAATCCCTTA | 42 | CORE |
| 34[62] | 33[48] | AGCCGATATCAAACCCTCAATCCTTGCT | 28 | CORE |
| 35[21] | 29[27] | AGGCTCCTTGGCAAAATGAAAGTTTTCAGTGAAATAACGGTAGAGCGGG | 49 | CORE |
| 55[28] | 43[41] | AGGGGGTAAAAACCTAGTAAGAGCAACAATTTAGGATTTGTA | 42 | CORE |
| 47[42] | 49[48] | AGTAAATTGATTAAGATAAGGCTTGAGAAAGCTAA | 35 | CORE |
| 16[83] | 6[77] | AGTACATCTGTTTAATAAAGTTCTTACCCAAGATT | 35 | CORE |
| 19[42] | 23[48] | AGTGAGAATGATGGCAGATAGAAAGGAAAGGAGCAAGAGGTGGTCCGTG | 49 | CORE |
| 6[76] | 7[69] | AGTTGCTGCAAGCCCAAGTACCGCACTCATCCACCCAGAACGAGAACAA | 49 | CORE |
| 18[41] | 7[41] | AGTTTTGATTAAAGCAGCATTCACCACC | 28 | CORE |
| 22[55] | 19[62] | ATAAAACCTAACAACTAATAGTGATTATCAATTCA | 35 | CORE |
| 0[125] | 3[118] | ATACATATTAAGACACAATGAAATAGCACGCTAAT | 35 | CORE |

| Start | End | Sequence | Bases | ID |
|---|---|---|---|---|
| 0[55] | 15[62] | ATAGAAAAAGAGAAGAGGGTTGAGCCACCAAATAA | 35 | CORE |
| 15[119] | 6[119] | ATAGCGAACATTTAAAGCCAAGTAATTTCAGTTACTAGCGAA | 42 | CORE |
| 20[69] | 24[63] | ATATTCCATTAGAGACCACCATTCACCAGAAATGG | 35 | CORE |
| 3[119] | 10[112] | ATCAGAGTTTTTGTTTAACGTAATTTGCAGGCAGA | 35 | CORE |
| 47[84] | 55[90] | ATCTACAAAACAGGAAGCAAATAGCCAGGAACAAAGCCAGCTTAAATAT | 49 | CORE |
| 22[118] | 19[125] | ATGCGCGTTTACAAACAATTCGTAACATTACCATA | 35 | CORE |
| 11[91] | 19[104] | ATGCGTTTTTTAATCGCGCAAAACATCGAGGATTTAGAAGTACAAAGAAATAATG | 56 | CORE |
| 16[104] | 7[104] | ATTACCTATACAAATCGAGCCAGTAATATGCAGAACGCGCCTCTTATCAATCATTA | 56 | CORE |
| 47[105] | 55[111] | ATTGCCTTGATAATACGTTAAAATTCGCTAATGGGCCTCAGGTCAAATG | 49 | CORE |
| 35[42] | 37[48] | ATTGTATCGACGTGACGGAGTTTATCAGCGCATAA | 35 | CORE |
| 51[119] | 47[125] | ATTTCGCAGGTGGCATCAATTGCAAGGATAAAAATAGATTCATGGAGCA | 49 | CORE |
| 6[69] | 1[69] | ATTTTGCAATTTTAAACAGGGGACGGGATAGCCGACAGAAGG | 42 | CORE |
| 52[97] | 45[104] | CAACATGGAGTAGATTTAGTTTAGTAGTATTGTAA | 35 | CORE |

| Start | End | Sequence | Bases | ID |
|---|---|---|---|---|
| 44[76] | 48[70] | CAACATTATAGGCTGGTGTATAAGATCTGACCTTCGCTATTTTGATATT | 49 | CORE |
| 48[69] | 54[63] | CAACCGTAAACATTCAAGGCAATAACAGACGGTGTTTTAATTGCCCGAA | 49 | CORE |
| 35[105] | 29[111] | CAAGAATTCATTGCGTCATAAGGTGCCCCTAACTCCCAACGCCCACGCT | 49 | CORE |
| 46[104] | 35[104] | CAGAAAATAAAAAAGTTTTTTGCAACCG | 28 | CORE |
| 20[91] | 24[84] | CAGAAGGATACATTTGCCATTAAATAAAACATTTTG | 36 | CORE |
| 42[55] | 55[55] | CAGGCCGAAAGAGGAAATGTTTAGACTG | 28 | CORE |
| 37[112] | 48[105] | CAGGCGGCCCCGGTGAGAGTCAAAGGGTGAGAAAG | 35 | CORE |
| 24[48] | 14[35] | CAGTTGAGTAAAAGCAGGGCGCGTACTAAATCAAGGTATAGC | 42 | CORE |
| 39[112] | 35[125] | CATGTTTCAAAAATTATTTTGTTAAAATATATGTACCTTTAGCCGTTCCGCAGCAC | 56 | CORE |
| 51[77] | 47[83] | CCAATTCATAAATCATACAGGATGACCCTGTAATAATCAATATTGAGAG | 49 | CORE |
| 15[28] | 17[41] | CCCTCAGCGGGTACTGCCTGTTCTTCGCAGGCGGTATTCCAC | 42 | CORE |
| 8[48] | 0[42] | CCGCCGCCCAGAATGTCTCTGGTCAGTGCAGTGCCACTCCTCATTCATA | 49 | CORE |
| 7[105] | 1[111] | CCGCGCCACTTGCGAGAGCCTCAAAAATAATTGAGATAGCTAAAGAACT | 49 | CORE |
| 6[118] | 8[105] | CCTCCCGCAATAGCATCGGCTGTCTTTC | 28 | CORE |

| Start | End | Sequence | Bases | ID |
|---|---|---|---|---|
| 11[49] | 9[62] | CCTTGGAATCATAAAAGTAATTCGCG CAGGAAATGTCCAGAC | 42 | CORE |
| 54[41] | 32[35] | CGACGATAATAGTACAAAAGAAGGCA CCTGAGGACCGCTCACGAATGCG | 49 | CORE |
| 26[125] | 12[119] | CGGGCGCAGCCGGCAATCATACAATC GCTTCTGAC | 35 | CORE |
| 27[112] | 24[119] | CGGGGAATAGGGCGTGCCTGAGTAGA AGCATTGCA | 35 | CORE |
| 38[48] | 41[55] | CGGTCGCGGGTAGCGAGGCTTAACCT AAAAAAAGC | 35 | CORE |
| 27[70] | 15[83] | CTAAATCCACACCCAACCGTTGTAGC AAACGCTCAATCGTCG | 42 | CORE |
| 12[118] | 17[118] | CTAAATTCAGTATAACAATTTATTTCA A | 28 | CORE |
| 7[42] | 1[48] | CTCAGAGATCAAAATATTAGCCACCG TATCACCAATTATCACACGGAAA | 49 | CORE |
| 43[49] | 44[49] | CTGATAAATGTAACAGCGATGTGTCG AA | 28 | CORE |
| 55[70] | 44[77] | CTGCGGAAGAGGAACGAGCTTCAAA GCGTATGCAATCCGTGGCTTTCAT | 49 | CORE |
| 33[63] | 29[69] | CTGTTGCGCCTGTGCGGAAGCTGGTT TTTGCCCTT | 35 | CORE |
| 55[112] | 43[125] | CTTTAAACTATTATTCAGGATTAGAGA GCTTAATTACGTTGG | 42 | CORE |
| 2[83] | 5[76] | GAAAAGTAAGCAGAGAATTAACTGA ACAGAGAATAACATAAATCCTGAA | 49 | CORE |
| 19[105] | 27[111] | GAAGGGTTCAGATGATTATTCCATTTG AGAATCCTGAGACTAAGCTTGA | 49 | CORE |

| Start | End | Sequence | Bases | ID |
|---|---|---|---|---|
| 1[28] | 14[28] | GAAGGTACCAGCGCATTAAGACCGGAAT | 28 | CORE |
| 36[27] | 28[21] | GAATTTCAATGACAAGTTAAACGAAAGATTTTTCATAATGCCAAAGGGC | 49 | CORE |
| 23[77] | 19[83] | GACCAGTAAAATACCGAACGACCGTCAATAGATAAGCGGAATATTGTTT | 49 | CORE |
| 9[63] | 0[56] | GACGACAAAAGGTATTACTAGATAAGAAGATTAGCTTTTGTCACAATCA | 49 | CORE |
| 47[63] | 55[69] | GAGGGTAATCAAGACAGGCGCAAATGTGAACCCGTCTGGTGCTCCAATA | 49 | CORE |
| 31[98] | 38[91] | GAGTGAGCCTGCATCAGACGAAGGTTTCGACGCAG | 35 | CORE |
| 16[62] | 18[49] | GAGTGCCCTCATTTCCTCATAGTTCGCCCGGATTAGCGTAAC | 42 | CORE |
| 55[56] | 40[49] | GATAGCGCGGAAACGCCATTCTTTCAGAGGCTACA | 35 | CORE |
| 18[48] | 27[55] | GATCTAAAGACAGCTCAGGGAACCCTCAGATATAATTTTTTGGGGTCGA | 49 | CORE |
| 4[97] | 8[91] | GCAGCCTACGAGCGTCTTTCCGGAGGTTTTGAAGCCGTAGGATTCCAAG | 49 | CORE |
| 7[21] | 1[27] | GCCACCCGCCACCACATTTTCAATCAAGCCATTAGGCCATTTAGGGAGG | 49 | CORE |
| 26[55] | 22[56] | GCCGCTAAGTCTGTCCATCACATTATTTACCCTCAAGCCTATTGGCAGAGCAGAAG | 56 | CORE |
| 20[27] | 26[21] | GCGAATAAAGGTTAAACACCGGCACGCGCGAGCTCTATAATCTTTGACG | 49 | CORE |

| Start | End | Sequence | Bases | ID |
|---|---|---|---|---|
| 32[48] | 33[62] | GCGGCCAAATTCCACACAACATACGAGCCACTCTGTGGTGCTGAACCTCAAGGTCA | 56 | CORE |
| 32[34] | 37[34] | GCGGGCCAATCTAATGCAGGGACAACCA | 28 | CORE |
| 36[111] | 28[105] | GCGGTCCAGCCGCACTCACGGACAGCGCCCAGGGTGGCCTCTTCAAAAG | 49 | CORE |
| 34[76] | 35[62] | GCGGTTGCGTCAGCCGGCCAGAGCACATCCTCATACCGGACT | 42 | CORE |
| 12[76] | 17[76] | GCGTTAAAAAAAGCAAATCAATTTGAAT | 28 | CORE |
| 19[84] | 14[77] | GGATTATGTACCTTTACAAAATGGAAACCTATTAAGGCTTAGGTAGTAC | 49 | CORE |
| 1[112] | 15[118] | GGCATGAAAGGTGGGCAAATCGGTCTGATGAAAAC | 35 | CORE |
| 13[35] | 10[28] | GGCTGAGCGTATAAGGAAACGATCAGTAGCGACAGGGTCATACGTTCCA | 49 | CORE |
| 49[91] | 47[104] | GGGAGAAGCCTTTAGCCGGAGTCAGGTC | 28 | CORE |
| 8[27] | 0[21] | GGTTGAGACAAATAGTAAGCGCTGGTAAATGCCCCTGAAAGTCAAAGAC | 49 | CORE |
| 29[112] | 51[118] | GGTTTGCTTATAAATCGCTATGTATCGGATAGGTCGCTGAATTAGATAC | 49 | CORE |
| 40[97] | 50[91] | GTCACGAAATCGGCCTTCCTGTATTTAAAGCATTA | 35 | CORE |
| 44[104] | 36[91] | GTCTGGCGAAACGTAAAAAGATTTGCTCAGGCGCTTTCGCACGTCAGCACGTCTCG | 56 | CORE |

| Start | End | Sequence | Bases | ID |
| --- | --- | --- | --- | --- |
| 26[104] | 4[98] | GTGTAGCGATTTAGCCTTTTTATGTAAATTGAAATAGAGGGTGAAAATA | 49 | CORE |
| 28[83] | 39[76] | GTGTTGTTGGCCCTATTGGGCGCCAGGGATAAAGTGGCCAGTACGGATA | 49 | CORE |
| 9[105] | 0[105] | GTTTATCGGCATTTTTCTTACTAATGGTTGCTGATCAACATA | 42 | CORE |
| 0[104] | 12[91] | TAAAAGATACCCAATCTTACCGAAGCCCCAAAGTCACCGACC | 42 | CORE |
| 40[125] | 38[112] | TAAGTTGTTGCGTTCAGCCAGCGGTGCCACATCCCGATAGCT | 42 | CORE |
| 11[28] | 19[41] | TAAGTTTCCCAATACTGTAGCCAGTATTTCTAAAATATCTTTCAACTAATCAGCGG | 56 | CORE |
| 25[84] | 27[90] | TACTTCTCTGCGCGTAACCACGGAACCCTAAAGGG | 35 | CORE |
| 16[41] | 8[28] | TAGCAAGTAACGGGAATTTACGCCCCCTTCACCGGAACCAGATCAGAGCGACAGGA | 56 | CORE |
| 48[132] | 54[126] | TAGGTAATTTTAGAGCTGAAAAAATGGTCTTAGAGTACCTTTGGTCTTT | 49 | CORE |
| 19[63] | 27[69] | TCAATATACAATAATGATTGCTATATGTCCTTGCTATAAGTGTAAAGCA | 49 | CORE |
| 27[28] | 15[41] | TCACCCATGGTTGCAGTGAGGCCACCGAGGATCCCAACCGCC | 42 | CORE |
| 55[91] | 43[104] | TCATTGACGGATTGCCGGAAGCAAACTCTGTAGCTTTGACCG | 42 | CORE |
| 28[76] | 51[76] | TCCAGTTAGGCTGCACCGCTTCGGATTCCTAAAGTTTGATTC | 42 | CORE |

| Start | End | Sequence | Bases | ID |
|---|---|---|---|---|
| 36[90] | 28[84] | TCGCTGGATAAAAAAAACAGCCGGAAACCGTTGTATTGGGAAGGGTTGA | 49 | CORE |
| 23[98] | 20[92] | TCTGGCCAGCCCTAGAGGCGAAATATACAGTAACAACTTCTGAACCAC | 48 | CORE |
| 3[49] | 5[62] | TGAAACCATCATTAAAGCGCGATAGCAGGTTTGCCATGCTAC | 42 | CORE |
| 46[55] | 43[48] | TGACAAGAACCGGAACAGATGATCCGCGTCATCGC | 35 | CORE |
| 13[77] | 1[83] | TGCTCTGGGTTATAAGACACCACGGAATGGAAACG | 35 | CORE |
| 28[41] | 40[28] | TGGACTCCAGGAGGGGATTTTAGACAGGTGTTATCTAAAGAC | 42 | CORE |
| 35[84] | 29[90] | TGGTCTGTCAATCCTGGGTAATCCAGCGCTGGGGTTTTGCGTGAGAGAG | 49 | CORE |
| 0[41] | 12[28] | TGGTTTAAATATTGCGTCACCGACTTGACAAGGCCACAGTTA | 42 | CORE |
| 50[48] | 54[42] | TTAAGCAAGGTAGAAGTTGAGCTATCATTACCAGA | 35 | CORE |
| 26[62] | 12[49] | TTAATGCGGTGCCGCCGTCGAGGATTAGTAAACACCGAGTAA | 42 | CORE |
| 17[119] | 7[125] | TTACCTGTTTCAGGGATAAGTATCAATAAAGCAAA | 35 | CORE |
| 1[49] | 2[49] | TTATTCATTGTTACACAAAAAAGGTGAA | 28 | CORE |
| 29[91] | 43[97] | TTGCAGCCGAGATAGGGCGATCACTCCACGGCGGA | 35 | CORE |
| 20[111] | 26[105] | TTGCGGAATTAGACAACTGATAACAGAGAAACGCTCATCACTCTGGCAA | 49 | CORE |

| Start | End | Sequence | Bases | ID |
|---|---|---|---|---|
| 49[105] | 54[105] | TTTCAACCTACTAATGACCATATAATGCCAACAGGAGTCAGA | 42 | CORE |
| 38[69] | 30[56] | TTTCTGCCACGGGAGCCAAGCGCCATTCTGGAACAAGAGTCCAGCTGATTCTTTTC | 56 | CORE |
| 7[84] | 1[90] | TTTTCATCTTAAATAACGCTATTACAGACCCTGAATTTTTAACAATAAT | 49 | CORE |
| 0[20] | 3[13] | AAAAGGGCCGATTGGGGAATTAGAGCCACAGTAGC | 35 | CORE |
| 51[14] | 45[20] | AAACGAATTGGGAAGAACCGA | 21 | CORE |
| 21[9] | 19[20] | AAGGAATTGAGGATAATTTGCTAAAC | 26 | CORE |
| 41[7] | 43[13] | ACGGGTATGACCCCACCAAGC | 21 | CORE |
| 26[20] | 13[20] | AGCACGTCACTACGTCACCGTTGAAACA | 28 | CORE |
| 29[28] | 51[34] | AGCTAAACAACGTCACTACGAATACACTAACGGAGAATACCAAACAACA | 49 | CORE |
| 34[20] | 36[7] | ATCAACAGTTGA??AATCTCCGCTTGAT | 28 | CORE |
| 16[20] | 7[13] | ATGTACCGAGTGTATCATACATCATCGGCCGGAACCGCCTCCCCACCCT | 49 | CORE |
| 13[7] | 24[9] | ATTATTCACTCAGGCCGCCACTAATC | 26 | CORE |
| 39[28] | 35[41] | CAGCATCCCATGTTAACTTTGAAAGAGGTATTCATTCGCCCACTTGCTTGCCTTTA | 56 | CORE |
| 45[7] | 47[20] | CATAAGGGAAAAATGCGATTTTAAGAACCATTGTGTGCCCTG | 42 | CORE |
| 54[20] | 41[20] | CGAGAGGAGAAGTTTCATCTTAAATACG | 28 | CORE |

| Start | End | Sequence | Bases | ID |
|---|---|---|---|---|
| 44[20] | 33[20] | CGGAACGTCAGCAGGGCCGCTAGCAGCA | 28 | CORE |
| 12[20] | 5[13] | CTGCCTAACCATTATTTGCCTTTAGCGTCGCGTTT | 35 | CORE |
| 9[7] | 20[9] | CTTGATAAATGAATGGATTTTTTTCA | 26 | CORE |
| 28[20] | 30[9] | GAAAAACAGAATCACGCCAGAATCCT | 26 | CORE |
| 25[9] | 15[20] | GAGAAGTGTTTTGAATTCGCCTCAGA | 26 | CORE |
| 33[9] | 48[7] | GAGCCTTTGCGGATAGTTGGCTGCTCTAAGGCTAATTACC | 40 | CORE |
| 11[7] | 22[9] | GATACAGGTAACACCACCAGTCAGTG | 26 | CORE |
| 31[9] | 44[7] | GCTGTGTTTCCAGTCACCCAGGCGCA | 26 | CORE |
| 53[14] | 55[27] | GGAATTACTAATGCGCGAAACAAAGTACAAAACACTTGCCAG | 42 | CORE |
| 23[9] | 17[20] | GGGTTTCTGCCACCTGCAAACAAACT | 26 | CORE |
| 18[20] | 7[20] | GTTAGTATTCACAAGCAGGTCCAGAACC | 28 | CORE |
| 46[20] | 35[20] | TAACAAACGCCGACTTAAACAAAAAAAA | 28 | CORE |
| 46[34] | 48[21] | TACCCAAACACCAGATTTCAACTTTAAT | 28 | CORE |
| 40[20] | 32[9] | TGAGGAATTCCTGTCGGTCATACCGG | 26 | CORE |
| 49[21] | 54[21] | TGGCTCACAGGACGCTAACGGCATTCAACGAGGCAAAAATAG | 42 | CORE |
| 51[35] | 47[41] | TTATTACATAAAGCCTCCAGTTTATACAGAGCATATGGTTTAAACGAGT | 49 | CORE |
| 4[139] | 10[133] | AAATAAGACAGCCAAACAACG | 21 | NEIGH |

| Start | End | Sequence | Bases | ID |
|---|---|---|---|---|
| 6[139] | 8[133] | AACGCGATAGAAGGGCATGTA | 21 | NEIGH |
| 9[133] | 20[133] | AAGAAAAGAAATTGTATTTGCTTTTAAA | 28 | NEIGH |
| 2[139] | 12[133] | AATAAGACCACAAGTAGTTAA | 21 | NEIGH |
| 13[133] | 23[139] | AGAACGCAGTGAATGATTAAGATATTACAGCGTAA | 35 | NEIGH |
| 55[133] | 53[139] | AGAATGAAAAATCAAATTGCT | 21 | NEIGH |
| 20[132] | 26[126] | AGTTTGAGACAACTGCTATTAACCTGAACGCCAGCAACTCAAAAAGGAG | 49 | NEIGH |
| 11[133] | 21[139] | CAGTAGGAAACAAAGAAGATGTTGAATGCGTATTA | 35 | NEIGH |
| 42[139] | 51[139] | CAGTTTGGGGCGCACGGATGGCAATAAC | 28 | NEIGH |
| 35[126] | 29[132] | CGTCGGTAGGTGTCGGTGTGTTTACCTGGCGCTCACCAGCTGGGCGAAA | 49 | NEIGH |
| 46[139] | 48[133] | CTAGCATGAATCGATAATGTG | 21 | NEIGH |
| 36[132] | 28[126] | CTGATTGTGATGAACTCCGTGCCGGAATAGGCGATGCTGGCGATCGGCA | 49 | NEIGH |
| 8[132] | 0[126] | GAAACCACCTGAACCCAACATCGCTCAATTTCATCAAGACAAAAAATAC | 49 | NEIGH |
| 19[126] | 27[132] | TCAAAATCGTAGATAGCAAAAATTAATTTAGCTTATTATCAAGAACGTG | 49 | NEIGH |
| 7[126] | 1[132] | TCAGATAGGCGTTTAAAATAAAAACGATAGATAACGCAAGAATCCTTAT | 49 | NEIGH |
| 44[139] | 49[139] | TTTAACCAAATTTTGGCGCGAACCCTCA | 28 | NEIGH |

**Table S2B. List of Core and Neighbour Strands (H2)**
This table denotes the ssDNA core and neighbour staples sequences used for the H2 hinge. The Start and End columns denote the helix[base] positions at the 5' and 3' ends of the DNA staple strands in the CaDNAno design for the respective designs (Supplementary Fig. 1B). Refer to Table 1 to obtain definitions and colors of the IDs.

| Start | End | Sequence | Bases | ID |
|---|---|---|---|---|
| 0[104] | 3[97] | AACATGTAAAGCGAATTATAGTCAGAAGCAAAGAA | 35 | CORE |
| 0[125] | 1[111] | CTGAATAAAACTCCAATCAGGTCTTTACCCTGACTACCAGAC | 42 | CORE |
| 0[146] | 12[133] | GGATGGCAGAGAGTCGAGAATGACCATAATTGAATTGTACAG | 42 | CORE |
| 0[62] | 12[49] | TTCATTCAAGCCCGAGGAATAACCTGTTAATAGTAAGCCAGC | 42 | CORE |
| 0[83] | 12[70] | TAAAGTATCGCGTTGATTGCATCAAAAACAATAAAGCCTCAG | 42 | CORE |
| 1[112] | 14[112] | CGGAAGCTAATGCTAGCCGGATACAACG | 28 | CORE |
| 1[134] | 23[139] | AGGATTTTAGAGCGTCAATCTATACCAAGGCAAACCGAGTAGAAATAC | 48 | CORE |
| 1[70] | 24[70] | TCAAATACGGTGTCGGGCCTCCGCCAGCAAGTTTCATAACAT | 42 | CORE |
| 10[132] | 0[126] | GGTTTAACTCATTCACCAGGCGCAGACGTTAATTG | 35 | CORE |
| 11[105] | 0[105] | CAAATCAACCTTCAGTTACTTGTAGCTC | 28 | CORE |
| 11[112] | 22[112] | ACGTAACAGCGAAAAACCATCGTTGCCC | 28 | CORE |
| 11[133] | 21[139] | AGTGAATTGCGGGAGATATATTGCGGTATCGTCTC | 35 | CORE |

| Start | End | Sequence | Bases | ID |
|---|---|---|---|---|
| 11[147] | 0[147] | GCCCTGAGGACAGAAACCGAATTTTTGC | 28 | CORE |
| 11[56] | 7[55] | CCGTGCATAACAACTATATTTGCCGGAGACAGTCAGAGGGTA | 42 | CORE |
| 11[70] | 21[76] | GTTTGAGGGCTTTGTTTACCACATAAACTTAGTGA | 35 | CORE |
| 12[104] | 3[90] | TCAAGAGTTAGCAATAAAGCCTCAGAGCATAAAGCGGCAAGG | 42 | CORE |
| 12[146] | 10[133] | TGAACGGCCCCCTCGTAATAGTAAAATGAGCGAGATTGAGAT | 42 | CORE |
| 12[48] | 0[42] | TTTCCGGCGCAACTCAGTTGA | 21 | CORE |
| 12[97] | 15[90] | TAATCTTGGATATTGAGGGTAGCAACGGCGGGTAA | 35 | CORE |
| 13[56] | 4[56] | AGGGCGAGCACTCCGTAGCATATTATGA | 28 | CORE |
| 14[90] | 0[84] | TAAATTGTGTATTATTCGCTCGAAATCCATGCAAC | 35 | CORE |
| 15[112] | 17[125] | GGCACCATCACGCAGCAACAGGAAAAACGGTCACTGCCCACG | 42 | CORE |
| 15[49] | 27[69] | AACGACGGCTTTTTCATGAGGTGGCGAACCCGAGA | 35 | CORE |
| 16[104] | 6[98] | TCGGAACCATTACCATTACCTACCCTCGGCATAGT | 35 | CORE |
| 16[125] | 6[119] | CCTCAGCAAAGCTGTTTCAACATAAAAAACGCCA | 35 | CORE |
| 16[146] | 6[139] | CCGCTTTAAGGCTTATTGGGCGGCTTTTATTCAACT | 36 | CORE |

| Start | End | Sequence | Bases | ID |
| --- | --- | --- | --- | --- |
| 16[152] | 27[157] | TAAAGGACTAAAATGACCCCACCGTCTATCA | 31 | CORE |
| 16[48] | 25[55] | GCTTTCAGTTGTAACAAGGCGCCTTATAAATCAAAGCTGGTTCTCTGTG | 49 | CORE |
| 16[83] | 3[83] | CTACAGAGGGACGACGGAACCGACAAACAGTATCGTCATACA | 42 | CORE |
| 18[104] | 7[104] | AAACAGCTTATACCAGCCCCAGAGAGTC | 28 | CORE |
| 18[125] | 7[125] | TGCTTTCGAAGAAAACCCCGGGAATCGA | 28 | CORE |
| 18[62] | 7[62] | AAGGGATTTCGCGTGTAAACGGCTATTT | 28 | CORE |
| 18[83] | 10[77] | AGATAGAGCCAGCTTTAGAACTTTTACATCAACAT | 35 | CORE |
| 19[105] | 27[111] | GGAATTGATTTCTTTGACAACGACAGCACTACGAAGAGATTTCCACTAT | 49 | CORE |
| 19[125] | 27[132] | TTTCACGTATCAGCTCATAACCTCGTCACACGAAAGAGCGCGATCCAACG | 50 | CORE |
| 19[63] | 22[49] | TCAACAGCGTGGTGCGCCATGAGGACTAAAGACCAGTGCCAAGGCGAAATGTGTTC | 56 | CORE |
| 2[158] | 1[165] | CAGTTCAGAAAAACCTTTAATTGCTCCTTTT | 31 | CORE |
| 2[90] | 24[91] | CAAAGCGTTAATTCGAGCTTCTTTAAATGCGACCTCGCCTGAAATACGTGCAATAC | 56 | CORE |
| 20[111] | 26[105] | ACGATCTGCCGTTCTGCGGCTAGCCATTAATTAACTTCTGCCAACAGCT | 49 | CORE |

| Start | End | Sequence | Bases | ID |
| --- | --- | --- | --- | --- |
| 20[131] | 26[126] | CAGCCCCGTTTTTGAGCCGGCTCATGAAAGAGTTTCTTCGCTTTTCA | 48 | CORE |
| 20[160] | 22[147] | AAACTACAACGCCTGCAGCCTCCGCCGG | 28 | CORE |
| 20[69] | 26[63] | ATTTTCTGCGGCCTATCCCTTACTATCGCACTTGCGGCCAGAAGCAAGC | 49 | CORE |
| 20[90] | 24[84] | TTTCCAGTAAAGTTGTTTCTTATCCAGATTCTTTG | 35 | CORE |
| 21[112] | 19[124] | CGGCAAACGCGGTCTCATAGTTAATTT | 27 | CORE |
| 21[140] | 17[152] | GTCGCTGGTAGCATCTCCAAAAAAAAGGAGCCTTTCTGAGG | 41 | CORE |
| 21[77] | 19[90] | TGAAGGGACGTTAGGGAGTGAGAATAGA | 28 | CORE |
| 21[91] | 19[104] | AAACGATGCTGATTAAAGTTTAACTAAA | 28 | CORE |
| 22[160] | 21[165] | ACTCAATCCGGCCAGAGCA | 19 | CORE |
| 23[140] | 10[147] | CTACATTGCGCGGTTCGGTCGAATTGTAAAACGAACTAACGGGTAGTAA | 49 | CORE |
| 23[147] | 26[147] | TTGACGCAATCAGTCTCACAGTTGGGCG | 28 | CORE |
| 23[48] | 17[62] | GGTGCCAGAACTCAAACACTGGCGTACAG | 29 | CORE |
| 23[70] | 17[83] | GCCTTGCTGGTAATTGCTCGTGTCCCGG | 28 | CORE |
| 23[91] | 17[104] | ACAATATTACCGCCGGTAATGCCGACAA | 28 | CORE |
| 24[160] | 23[165] | TTTTTATTCAATCGTCTGA | 19 | CORE |

| Start | End | Sequence | Bases | ID |
|---|---|---|---|---|
| 25[91] | 15[104] | GTCATACCGGGGGTCGTTGTAAATGCCA | 28 | CORE |
| 26[104] | 13[104] | GATTGCCCAAGAGTGTATCATGCTCCAT | 28 | CORE |
| 26[125] | 4[118] | CCAGTGACGTGGACAACAAAGACGAGGCGCATAGGCGTCATATGGATAGC | 50 | CORE |
| 26[146] | 13[146] | CCAGGGTGCGAAAACAGCGATATAAGGG | 28 | CORE |
| 26[160] | 25[165] | TTGCGTATTGAGGATCCCC | 19 | CORE |
| 26[62] | 14[49] | GGTCCACAGAATAGAGGGGGATGTGCTG | 28 | CORE |
| 26[83] | 26[91] | CCTGAGAGAGTGTTGTTCCAGTTTGGAACTTCACC | 35 | CORE |
| 27[112] | 15[125] | TAAAGAAGACGGGCAGCACGCGTGCCTGCTGTCCAACCTAAA | 42 | CORE |
| 27[133] | 15[146] | TCAAAGGGGTTTTTCGTCCGTGAGCCTCGAGGCCAAGAATAC | 42 | CORE |
| 27[70] | 15[83] | TAGGGTTGAGTTGCATGCGGCGGGCCGTATTAGTACATTAAA | 42 | CORE |
| 28[104] | 40[91] | GGTGCCGCAGTCGGGCTCACAATTCCACGCGGGAGAGAGCCG | 42 | CORE |
| 28[125] | 40[112] | AATCAAGCAGCTGCTCCTGTGTGAAATTGATTAAAACCCTCA | 42 | CORE |
| 28[146] | 40[133] | CACTACGCAACGCGGTAATCATGGTCATGAACGGTGCCGCCA | 42 | CORE |
| 28[62] | 30[55] | GATTTAGAGCTAACTAAAGCCT | 22 | CORE |
| 29[112] | 51[118] | GTCGTGCTTTTTTGTTATTAGTTAGCAATTATTCAGAGCATGTACCGCG | 49 | CORE |

| Start | End | Sequence | Bases | ID |
|---|---|---|---|---|
| 29[133] | 42[133] | AATCGGCTGAACCATCATAATTCACCAG | 28 | CORE |
| 29[70] | 42[70] | AATTGCGTAAAGGGGACTGTAGCACCGT | 28 | CORE |
| 30[165] | 31[160] | GGGTACCGAGCTAGAAGTG | 19 | CORE |
| 30[54] | 28[42] | GGGGTGCCGCGGTGTGTATAATGAGTGAGCTTGACGGGGAA | 41 | CORE |
| 30[76] | 44[63] | GGAAGCAGCTTTCCACCAGAGGTCAGACCAAAAGGGCATTAT | 42 | CORE |
| 32[146] | 38[133] | CAGATTCACCAGTCTCAGCGGCAGGAGT | 28 | CORE |
| 32[153] | 29[160] | ACATTGGAATCCTGCGAATTCCGGGGAGAGGCGGT | 35 | CORE |
| 32[165] | 33[160] | AATGGATTATTTCTTTCGC | 19 | CORE |
| 32[55] | 39[47] | CACAGACCGAACCAATTTGAGGATTTAGAACGTT | 34 | CORE |
| 32[69] | 29[69] | GTAAGAAATAACGTTAAAGTGTCACATT | 28 | CORE |
| 32[76] | 36[70] | TGAAAGCAACAGAGGTCACCGGGCAGGAGGCGGTCCCGCCACTCAGGAG | 49 | CORE |
| 33[56] | 34[49] | CCAGCAGACACCGCCTGCAAC | 21 | CORE |
| 33[84] | 29[90] | TCGGTGGGAACCCTAATCAGAACAACATCACTGCC | 35 | CORE |
| 34[125] | 36[112] | CGTGGTGCTGGTCTAGCCCAAGGGTTGA | 28 | CORE |
| 34[165] | 35[160] | CATCCTCATAACCCAGTAC | 19 | CORE |

| Start | End | Sequence | Bases | ID |
|---|---|---|---|---|
| 34[48] | 32[56] | AGTGCCATACCGAAAATATTTTGGTTGCTTTGACGAGCACGTTACGTGG | 49 | CORE |
| 35[106] | 29[111] | ATAGCAGGTCAGCACCAGCTACATTCTGGAGGCCGTTATCCGAAACCT | 48 | CORE |
| 35[127] | 29[132] | CATGTAACGTCAGTCCAGCAACACGACTAGACAGAGCTGTTATTAATG | 48 | CORE |
| 35[147] | 36[133] | TTCGTCAGGAACGTGCCGGACTTGTAGACCGTAACGGCGGAT | 42 | CORE |
| 35[42] | 33[69] | AAACCCTCAAGAACCGCCACCCTCAGAAAGTATTAAAGATAA | 42 | CORE |
| 36[111] | 28[105] | TATAAGTTTATTCTGGGGTCAAATCCTCGAGCCACAGCCCCCGGGTCGA | 49 | CORE |
| 36[132] | 28[126] | AAGTGCCTTAAGAGGTACTGGGGAAAGCCCCTCAGCATCTTTTCACCCA | 49 | CORE |
| 36[162] | 38[147] | TAGCGGGGTTTTGCTCAGAGAAGGGCTTTT | 30 | CORE |
| 36[69] | 28[63] | GTTTAGTCTAATAGTCAATAGGAGGCAGCCGCCGCAGCGTCAAGCCCCC | 49 | CORE |
| 36[97] | 38[84] | GGAATAGGTGTATCCCCTGCCAGTGCCC | 28 | CORE |
| 37[112] | 48[113] | GAAACATGATAGCCGAATTGATGAAAA | 27 | CORE |
| 37[133] | 48[133] | GCTGAGACCTTTTTTAAGAGCTTTTTTG | 28 | CORE |
| 37[49] | 35[41] | AGGAGCAGAATACCAAGTTACAAAATCGTCTAAAATGGTCAGAAATATC | 49 | CORE |
| 37[70] | 35[83] | ATTAGGTTAATGCCACCGTACCCTCAGA | 28 | CORE |

| Start | End | Sequence | Bases | ID |
|---|---|---|---|---|
| 37[91] | 48[91] | TATTTCGGAAGGAATATCAGAAGAGAAT | 28 | CORE |
| 38[104] | 35[97] | GTGCCTTCCACGCAAGCAACCGCAAGAACCCTCAT | 35 | CORE |
| 39[112] | 49[119] | ATTAAAGCACAATCGTATGTTTTTGAAGTCTTTCC | 36 | CORE |
| 39[133] | 49[139] | GCAGTCTCACGGAAATACATAAACCTCCCCAGTTA | 35 | CORE |
| 39[147] | 28[147] | TACCGTTCCGCCTCACCGGAAGATGGCC | 28 | CORE |
| 39[48] | 38[56] | ATTAATCAGGAGGTTATAATAC | 22 | CORE |
| 39[84] | 30[77] | CTTGATACCACCAGTTCATCGCTAAATCGGAACCCTTGCGCTACGAGCC | 49 | CORE |
| 39[91] | 49[98] | TTCACAAATGGTTTATTAAGATTAGTTGATCTTACC | 36 | CORE |
| 4[117] | 12[112] | GTCCAATAAGCAAAATTACTGCGGAATCTGGCTG | 34 | CORE |
| 4[55] | 11[55] | CCCTGTAACCCTCACCGTCGGATTCTCCTGGGCGCATCGTAA | 42 | CORE |
| 4[83] | 10[70] | TAAATCGGTTGTACAATGCCTTAAATGT | 28 | CORE |
| 40[104] | 38[91] | CACCCTCCTAAACAGGCCAACAGAGATATGCCATCGAGTAAC | 42 | CORE |
| 40[125] | 38[112] | AACCGCCGGGATTTCAGTAATAAAGGGTACGGCTTTTTAAC | 42 | CORE |
| 41[56] | 41[55] | TGCCTTTCAGCATTGATTTAAAGTTTGTCATCATATAAGTT | 42 | CORE |

| Start | End | Sequence | Bases | ID |
|---|---|---|---|---|
| 41[91] | 51[97] | GCATTTTCCAATGAGAAGGTATAATCGGTATTTTC | 35 | CORE |
| 42[55] | 45[55] | GAATCTCTAGAACCTACCATATTGCACGTAAAACAAGAAACA | 42 | CORE |
| 42[69] | 53[76] | AATCAGTAGCCTGTTTAGTATTTTTCACGCAAGA | 35 | CORE |
| 43[133] | 45[146] | TCACCGTTAATATCAATCAGATATAGAATTTAGCGCATAAAG | 42 | CORE |
| 43[42] | 54[49] | GAAGGGTCTGATTAGGAATCATAATTACAATTTCA | 35 | CORE |
| 43[63] | 51[76] | TCAAAGACATTCAACATTCCACGAGAAC | 28 | CORE |
| 44[125] | 33[125] | ATTTTGTCCAGAATTAATAAGGGAGGTG | 28 | CORE |
| 44[132] | 55[132] | TAAGTTTTGAATTATAGCACCAGGGCTT | 28 | CORE |
| 44[146] | 33[153] | AAGACACCTGAATTGATGATAGGTCATTGCAGGCG | 35 | CORE |
| 44[158] | 43[158] | AAAAGAAACGCATTGAGCCATTTG | 24 | CORE |
| 44[48] | 38[35] | GAAATAAAGAAATTTTGCCCGAAGTATT | 28 | CORE |
| 45[105] | 55[112] | TTACGCAAATAGAAACGGAAAGGCCGGAATAAAGCC | 36 | CORE |
| 45[56] | 37[48] | ATAACGGTTGCTTTCTAACAAACCGCCACCCTCATCAATATCTATCTTT | 49 | CORE |
| 46[146] | 35[146] | CCGAAGCCTCCTCAAGTACCAACTGAGT | 28 | CORE |
| 46[158] | 45[158] | ATAGCTATCTTAGTGGCAACATAT | 24 | CORE |

| Start | End | Sequence | Bases | ID |
|---|---|---|---|---|
| 46[76] | 46[77] | ATAATAACGCCTGAATTCGGAATACCCAAAAGAACAAACGCA | 42 | CORE |
| 47[63] | 50[63] | ACAAAGTCAGAGGGGAAGCGCATTAGACTGAATAAGTAAATC | 42 | CORE |
| 47[84] | 42[84] | GCGCTAAACCGAGGTGGCATGACCAGCGAGGGAGGAACCATC | 42 | CORE |
| 48[112] | 54[105] | TAGCAGCCAACGAGCGCCTTAAGAATCATTAGAAACGACGACGTAAGAGA | 50 | CORE |
| 48[132] | 54[126] | TTTAACGTAATTTGCGACTTGGCAAGCACCATCCTGTTCAGCGAGGCAT | 49 | CORE |
| 48[162] | 50[147] | ATCCCAATCCAAATAAAACAGCCGAGGCGT | 30 | CORE |
| 48[90] | 52[84] | AACATAAATCCTGACTATTTTCCGTTTTCTGTCTT | 35 | CORE |
| 49[120] | 45[132] | AGAGCCTCAAAAAGTTAAGCCCAATAAAAGAAAAGTAGAAA | 41 | CORE |
| 49[140] | 47[158] | CAAAATAGAAACGAAAGAAACAATGAAATAGCA | 33 | CORE |
| 49[36] | 48[42] | AACAGTATTAATTATGAAACAAACATCAATGAACTTAACGAAAACAAA | 48 | CORE |
| 49[42] | 50[56] | ACATAAAATTTTCCTCAATAGTGAATCAAGTACCGGTCGCTA | 42 | CORE |
| 49[99] | 46[105] | AACGCTTTTACAGGAGATAACCCACAAGAACAAA | 34 | CORE |
| 5[42] | 13[55] | TTTTAGAATACTTTTTCTACTTAGCTATTTTCGCAAATGGTCCATATAAGTTGGGA | 56 | CORE |
| 5[77] | 8[71] | GAGTAATAGATTCAAAAGGGTATCTACACAAATA | 34 | CORE |

| Start | End | Sequence | Bases | ID |
|---|---|---|---|---|
| 50[55] | 47[62] | TTAATTATCAATATATGTGAGGGGAGAAACCCTGA | 35 | CORE |
| 51[119] | 35[126] | CCCAATACGGGAGGAGCAAACGTAAGCAGAAAGTAGTCGAGATAGGAACC | 50 | CORE |
| 51[147] | 54[146] | GGCTTATCTGAACACCTGTTTCGCCAACA | 29 | CORE |
| 51[35] | 45[48] | AGAAGAGCTTAGAACCTTTTACATCGGG | 28 | CORE |
| 51[63] | 54[62] | CACTCATAGAACGGATCCAATAATATATT | 29 | CORE |
| 51[77] | 47[83] | AAGCAAGGCACCCAGCCTTCTCCTTGTACAATTTTAAACAGGTAATTGA | 49 | CORE |
| 51[98] | 35[105] | ATCGTAGATCAAGACTCCTTAGTTACCAGAACCTAATAGCCCTTTCAGGG | 50 | CORE |
| 52[83] | 38[70] | TCCTTATCCGATTGCCAAAGAGATTGGCGTATAAACAAGCCG | 42 | CORE |
| 53[49] | 54[42] | AAATGCTGATGCAAGTATTAAACTTATCAAAATCATATATGTTCTTCTG | 49 | CORE |
| 53[77] | 54[77] | CAAAGAACGGGTAACAAAACGAGAAAAC | 28 | CORE |
| 53[98] | 39[104] | CTGTCCACAATCAAAATATTGAATTCATACAAATA | 35 | CORE |
| 54[104] | 41[104] | ATATAAATACCAGTAACGTCACGGTCAT | 28 | CORE |
| 54[125] | 41[125] | TTTCGAGCAACAGTATTACCACGTTTGC | 28 | CORE |
| 54[145] | 31[146] | TGTAATAATCGCCAGCAAAACAAAATCCCTCAGAACGCCAG | 41 | CORE |

| Start | End | Sequence | Bases | ID |
|---|---|---|---|---|
| 54[41] | 31[41] | ACCTAAAAAACACCTCAGATGGGAATTAAGTAACACGTACTA | 42 | CORE |
| 54[61] | 42[56] | TTAGTTTAGAAAAAGCGACA | 20 | CORE |
| 55[113] | 43[125] | AACGCTCCAGTAAACAATAAACAACATAATTTACTTAAAGG | 41 | CORE |
| 55[133] | 43[146] | AATTGAGTTAGGCATAATGCAGAACGCGAGAAAAACACCGAC | 42 | CORE |
| 55[77] | 32[77] | CATATGCGATAGCAGCGCGTTAACCACCTCGTTAGTCTGACC | 42 | CORE |
| 55[84] | 53[97] | GTTATACAAATTCTGTACCGAAGTAATT | 28 | CORE |
| 6[118] | 10[105] | AAAGGAAAACAAGATTGATAATCAGAAAAGTCAGGATTGTGA | 42 | CORE |
| 6[138] | 9[132] | AATGCATAATCGTTGTCAATCATATGTAATCTAC | 34 | CORE |
| 6[97] | 10[91] | AAGAGCAACGGTAAGTGTAACTATCATATATGCGA | 35 | CORE |
| 7[105] | 10[112] | TGGAGCATTACGAGTTTACCAGACGACGTTTAATC | 35 | CORE |
| 7[126] | 1[133] | TGAACGGGATACATCCAAAATTTTAGACAATATTCAATCAAAAACAGGTC | 50 | CORE |
| 7[63] | 1[69] | TTGAGAGGAGAAAGTAAATGCCAAAAACTAACATCGATTAAGAAAGACT | 49 | CORE |
| 7[85] | 7[84] | CAGGTCATTGCCTAAAACAGGAAGATTGTATAAGAAGGCTAT | 42 | CORE |
| 8[162] | 7[158] | ATTACAGGTAGAAAGATTTAGCAAAAACCATCAGTTGAGATT | 42 | CORE |

| Start | End | Sequence | Bases | ID |
|---|---|---|---|---|
| 8[55] | 17[48] | TTAATATTTTGTTAGCCATCAAAAATAAAGCTCTCACGGAAAAACAATC | 49 | CORE |
| 8[70] | 0[63] | TTTAAATTCTGGCCTGAGCGAGTCTGCCAGAAGATCTCGGTGCTGGAAGT | 50 | CORE |
| 9[112] | 20[112] | ACGTTGGGAGGTGACGAATAATAGCGTA | 28 | CORE |
| 9[133] | 20[132] | GTTAATATCGGTTTTGAAAATTCCACAGA | 29 | CORE |
| 9[70] | 20[70] | TCCTGTACTTTCTCTTTCAGCTAAATGA | 28 | CORE |
| 9[91] | 22[91] | TGGCTCATTGATACCGGTGAGAATTTATAGTTGCGGGTAAAG | 42 | CORE |
| 0[41] | 3[34] | TTCCCAAAGATACAATTTTCATTTGGGGAAAGGTG | 35 | NEIGH |
| 12[34] | 5[41] | TCTGGTGCACGTTGGTGTAGAGTGGGAATAAAAAT | 35 | NEIGH |
| 12[41] | 9[34] | CACCGCTGCATCAATGCGGGAGAAGCCTGCAAGGACAAACGGTAACCAA | 49 | NEIGH |
| 17[35] | 7[41] | CACCGGAAAGAGACTAGGAACAAATTCGAAATTAA | 35 | NEIGH |
| 18[34] | 19[62] | GCAGAAAAACTTAAATTTCTGCTCATTTGCGCTAAACAACTT | 42 | NEIGH |
| 21[49] | 23[47] | AAA AAG CCG CAC AGG TAT GGG ATT TTC GCC AGC AGC CCG TAA AGC AAA TGC CAG C | 55 | NEIGH |
| 25[56] | 26[42] | GTGCTGCCTGAGTAGAGGTGCCCCCTGCCTGTGCATGCCCCA | 42 | NEIGH |
| 26[41] | 13[41] | GCAGGCGCAAAATCATTAAGTCAGGCTG | 28 | NEIGH |

| Start | End | Sequence | Bases | ID |
| --- | --- | --- | --- | --- |
| 28[41] | 40[28] | AGCCGGCCTGGCAACACGCTGCGCGTAACAGGGCGTTATCAT | 42 | NEIGH |
| 29[27] | 51[34] | CTAGGGCGGAACGTGAAGGAGCATGGCAAACTTCTGGAGAGACGACGCTG | 50 | NEIGH |
| 32[41] | 35[34] | TTGAATGGCTATTAATCGCCATTAAAAACGCTGAGAGCCAGCGAACCTC | 49 | NEIGH |
| 33[21] | 29[26] | CTAAAACGTCTTTAGCCGCTACCACCACCGGGCG | 34 | NEIGH |
| 38[34] | 47[34] | AGACTTTAAGGTTACGCAGAGGAAGATG | 28 | NEIGH |
| 39[28] | 49[35] | AAATCCTGCGTAGAAACAGTATCCTTGATTAATGGA | 36 | NEIGH |
| 45[21] | 55[34] | ATACAGTTTTTCAGGGATTATTTCATCACGTTAAATAAGAAT | 42 | NEIGH |

**Table S3.** List of Polymerization Strands
This table denotes the polymerization staples sequences used for the H1 hinge. The Start and End columns denote the helix[base] positions at the 5' and 3' ends of the DNA staple strands in the CaDNAno design for the respective designs (Supplementary Fig. 1A). Refer to Table 1 for the definitions and colors of the IDs.

| Start | End | Sequence | Bases | ID |
|---|---|---|---|---|
| 7[14] | 8[6] | GGTGCCATCCCACGCAGTTCTAGCTGATCATTAAATTTTTGTTAAATTAAACGATG | 56 | POLY |
| 7[42] | 6[14] | TGCCGGAAATCACCATCAATATGATATTCAACCACCAGCTTACGGCTGG | 49 | POLY |
| 9[6] | 10[6] | GGGTAAAGTCAGCTCATTTTTCGGATTGACCGTATAGACTTT | 42 | POLY |
| 55[2] | 42[2] | GCGAGAAAGGAATGATAAATAAGGATATAATCCAGAGTCAAT | 42 | POLY |
| 53[6] | 54[3] | ACTATCGGCCTTGCTGGTACTCCGGCTTAGGGTTTGAAATACCGACCGTGGGGAAGAAAGCG | 62 | POLY |
| 51[6] | 52[6] | GAATACGTGGCAAGCTTAGATTAATACCTTTTTAACATATCCAGAACA | 48 | POLY |
| 5[14] | 4[13] | TCAGCAAATCGTTAACTCAACTTATTGGCATCAGATGCCGGG | 42 | POLY |
| 49[6] | 50[6] | AATCCTTTGCCCGAATTACCTTTTAAACATAGCGATCAGACAATATTT | 48 | POLY |
| 48[34] | 48[6] | ACATTTAACAATTTCATTTGAACGTTATTAA | 31 | POLY |
| 45[2] | 46[2] | ATGAAACAACAGATGAATATTCATTTCAGAAATAAA | 36 | POLY |
| 47[2] | 46[21] | ACGTAAAACAATTACCTGAGCAAAAGCGAATT | 32 | POLY |

| Start | End | Sequence | Bases | ID |
|---|---|---|---|---|
| 7[14] | 8[6] | GGTGCCATCCCACGCAGTTCTAGCTGATCATTAAATTTTTGTTAAATTAAACGATG | 56 | POLY |
| 43[2] | 44[2] | ACGCTGAGATGATTGTTTGTTTAACGTACATCAAGA | 36 | POLY |
| 41[7] | 28[0] | GAGAAAACTTTTAGAAACCACCAGGCGAGAAAGGAAGGGAAGATAGCAAACGTAG | 55 | POLY |
| 39[7] | 40[8] | GCTTAATTGAGAACAACTCGTATTTTTGCGGAACAATCAAATATATTT | 48 | POLY |
| 36[34] | 36[6] | TTGGCAAATCAACAGTTGACCTAATTTACGA | 31 | POLY |
| 37[6] | 38[6] | ATAATATCCCATAAGGAATTGAGGACAAACAATTCGATCGCCATATTT | 48 | POLY |
| 35[0] | 34[21] | CTTATCCGGAAAGCATCACCTTGCTAGCAAAT | 32 | POLY |
| 33[1] | 34[0] | TATTATTTACTGATAGCCGAAAAATCTTATTCTAAG | 36 | POLY |
| 29[0] | 30[0] | TACGCAGTATGTAAGCGAAAGGAGACCCGCCGCAAGCCCAAT | 42 | POLY |
| 31[0] | 32[0] | AATTGAGTTGCTTAATGCATGCGCGAATCCCAATCC | 36 | POLY |
| 27[13] | 26[13] | CCATAGTGGTTCCGAAATCGGAAAATCCTGTTTGATGAATCA | 42 | POLY |
| 3[13] | 2[14] | CTGCCCGCTTTCCAGTGCTGACGCGACGGGAAACCTGTCGTG | 42 | POLY |
| 25[19] | 24[19] | CCTTTTGATAAGCAGTGTCACTGCGCGCATCAGACGATCCAGCGAGGTCATTTTTG | 56 | POLY |

| Start | End | Sequence | Bases | ID |
|---|---|---|---|---|
| 7[14] | 8[6] | GGTGCCATCCCACGCAGTTCTAGCTGATCATTAAATTTTTGTTAAATTAAACGATG | 56 | POLY |
| 23[19] | 22[20] | CTGTTTAGCTATTGCCGGGTTACCTGCACGTTAACGGCATCAGAATTTTCATTTGG | 56 | POLY |
| 19[14] | 18[14] | TGAACGGTAGATCACAGCGATCGTAAAA | 28 | POLY |
| 17[13] | 16[14] | TGTTAAATCAACGGACGGGAGCTCATTT | 28 | POLY |
| 21[20] | 20[20] | TATATTTTAAATATCGACATAAAAAATTTGGGCGGTTGTGTACGCAATGCCTGAG | 56 | POLY |
| 15[13] | 17[34] | TCGTAACCGGGGTTTTCCCAGTCACGACGAGGTGGAGCCGCCATAACCT | 49 | POLY |
| 11[6] | 12[6] | TTGTGAGAGAATGGGATAGGTCCGGAAACCAGGTGTGCTGCA | 42 | POLY |
| 13[6] | 14[14] | AAAGGGGGACAAAGCGCCATTCGCCATTTGGGTAACGCCATGCATCTGC | 49 | POLY |
| 1[13] | 0[13] | ATCCTGTTTGATATTTAGTTTGACCATTTTCTGCGAACGAGTAGGGTGGTTCCGAA | 56 | POLY |
| 7[14] | 8[6] | GGTGCCATCCCACGCAGTTCTAGCTGATCATTAAATTTTTGTTAAATTAAACGATG | 56 | POLY |
| 7[42] | 6[14] | TGCCGGAAATCACCATCAATATGATATTCAACCACCAGCTTACGGCTGG | 49 | POLY |
| 9[6] | 10[6] | GGGTAAAGTCAGCTCATTTTTCGGATTGACCGTATAGACTTT | 42 | POLY |

| Start | End | Sequence | Bases | ID |
|---|---|---|---|---|
| 7[14] | 8[6] | GGTGCCATCCCACGCAGTTCTAGCTGATCATTAAATTTTTGTTAAATTAAACGATG | 56 | POLY |
| 55[2] | 42[2] | GCGAGAAAGGAATGATAAATAAGGATATAATCCAGAGTCAAT | 42 | POLY |
| 53[6] | 54[3] | ACTATCGGCCTTGCTGGTACTCCGGCTTAGGGTTTGAAATACCGACCGTGGGGAAGAAAGCG | 62 | POLY |
| 51[6] | 52[6] | GAATACGTGGCAAGCTTAGATTAATACCTTTTTAACATATCCAGAACA | 48 | POLY |
| 5[14] | 4[13] | TCAGCAAATCGTTAACTCAACTTATTGGCATCAGATGCCGGG | 42 | POLY |
| 49[6] | 50[6] | AATCCTTTGCCCGAATTACCTTTTAAACATAGCGATCAGACAATATTT | 48 | POLY |
| 48[34] | 48[6] | ACATTTAACAATTTCATTTGAACGTTATTAA | 31 | POLY |
| 45[2] | 46[2] | ATGAAACAACAGATGAATATTCATTTCAGAAATAAA | 36 | POLY |
| 47[2] | 46[21] | ACGTAAAACAATTACCTGAGCAAAAGCGAATT | 32 | POLY |
| 43[2] | 44[2] | ACGCTGAGATGATTGTTTGTTTAACGTACATCAAGA | 36 | POLY |
| 41[7] | 28[0] | GAGAAAACTTTTAGAAACCACCAGGCGAGAAAGGAAGGGAAGATAGCAAACGTAG | 55 | POLY |
| 39[7] | 40[8] | GCTTAATTGAGAACAACTCGTATTTTTGCGGAACAATCAAATATATTT | 48 | POLY |
| 36[34] | 36[6] | TTGGCAAATCAACAGTTGACCTAATTTACGA | 31 | POLY |

| Start | End | Sequence | Bases | ID |
|---|---|---|---|---|
| 7[14] | 8[6] | GGTGCCATCCCACGCAGTTCTAGCTGATCATTAAATTTTTGTTAAATTAAACGATG | 56 | POLY |
| 37[6] | 38[6] | ATAATATCCCATAAGGAATTGAGGACAAACAATTCGATCGCCATATTT | 48 | POLY |
| 35[0] | 34[21] | CTTATCCGGAAAGCATCACCTTGCTAGCAAAT | 32 | POLY |
| 33[1] | 34[0] | TATTATTTACTGATAGCCGAAAAATCTTATTCTAAG | 36 | POLY |
| 29[0] | 30[0] | TACGCAGTATGTAAGCGAAAGGAGACCCGCCGCAAGCCCAAT | 42 | POLY |
| 31[0] | 32[0] | AATTGAGTTGCTTAATGCATGCGCGAATCCCAATCC | 36 | POLY |
| 27[13] | 26[13] | CCATAGTGGTTCCGAAATCGGAAAATCCTGTTTGATGAATCA | 42 | POLY |
| 3[13] | 2[14] | CTGCCCGCTTTCCAGTGCTGACGCGACGGGAAACCTGTCGTG | 42 | POLY |
| 25[19] | 24[19] | CCTTTTGATAAGCAGTGTCACTGCGCGCATCAGACGATCCAGCGAGGTCATTTTTG | 56 | POLY |
| 23[19] | 22[20] | CTGTTTAGCTATTGCCGGGTTACCTGCACGTTAACGGCATCAGAATTTTCATTTGG | 56 | POLY |
| 19[14] | 18[14] | TGAACGGTAGATCACAGCGATCGTAAAA | 28 | POLY |
| 17[13] | 16[14] | TGTTAAATCAACGGACGGGAGCTCATTT | 28 | POLY |

| Start | End | Sequence | Bases | ID |
|---|---|---|---|---|
| 7[14] | 8[6] | GGTGCCATCCCACGCAGTTCTAGCTGATCATTAAATTTTTGTTAAATTAAACGATG | 56 | POLY |
| 21[20] | 20[20] | TATATTTTAAATATCGACATAAAAAATTTGGGCGGTTGTGTACGCAATGCCTGAG | 56 | POLY |
| 15[13] | 17[34] | TCGTAACCGGGGTTTTCCCAGTCACGACGAGGTGGAGCCGCCATAACCT | 49 | POLY |
| 11[6] | 12[6] | TTGTGAGAGAATGGGATAGGTCCGGAAACCAGGTGTGCTGCA | 42 | POLY |
| 13[6] | 14[14] | AAAGGGGGACAAAGCGCCATTCGCCATTTGGGTAACGCCATGCATCTGC | 49 | POLY |
| 1[13] | 0[13] | ATCCTGTTTGATATTTAGTTTGACCATTTTCTGCGAACGAGTAGGGTGGTTCCGAA | 56 | POLY |
| 7[14] | 8[6] | GGTGCCATCCCACGCAGTTCTAGCTGATCATTAAATTTTTGTTAAATTAAACGATG | 56 | POLY |
| 7[42] | 6[14] | TGCCGGAAATCACCATCAATATGATATTCAACCACCAGCTTACGGCTGG | 49 | POLY |
| 9[6] | 10[6] | GGGTAAAGTCAGCTCATTTTTCGGATTGACCGTATAGACTTT | 42 | POLY |
| 55[2] | 42[2] | GCGAGAAAGGAATGATAAATAAGGATATAATCCAGAGTCAAT | 42 | POLY |
| 53[6] | 54[3] | ACTATCGGCCTTGCTGGTACTCCGGCTTAGGGTTTGAAATACCGACCGTGGGGAAGAAAGCG | 62 | POLY |
| 51[6] | 52[6] | GAATACGTGGCAAGCTTAGATTAATACCTTTTTAACATATCCAGAACA | 48 | POLY |

| Start | End | Sequence | Bases | ID |
|---|---|---|---|---|
| 7[14] | 8[6] | GGTGCCATCCCACGCAGTTCTAGCTGATCATTAAATTTTTGTTAAATTAAACGATG | 56 | POLY |
| 5[14] | 4[13] | TCAGCAAATCGTTAACTCAACTTATTGGCATCAGATGCCGGG | 42 | POLY |
| 49[6] | 50[6] | AATCCTTTGCCCGAATTACCTTTTAAACATAGCGATCAGACAATATTT | 48 | POLY |
| 48[34] | 48[6] | ACATTTAACAATTTCATTTGAACGTTATTAA | 31 | POLY |
| 45[2] | 46[2] | ATGAAACAACAGATGAATATTCATTTCAGAAATAAA | 36 | POLY |
| 47[2] | 46[21] | ACGTAAAACAATTACCTGAGCAAAAGCGAATT | 32 | POLY |
| 43[2] | 44[2] | ACGCTGAGATGATTGTTTGTTTAACGTACATCAAGA | 36 | POLY |
| 41[7] | 28[0] | GAGAAAACTTTTAGAAACCACCAGGCGAGAAAGGAAGGGAAGATAGCAAACGTAG | 55 | POLY |
| 39[7] | 40[8] | GCTTAATTGAGAACAACTCGTATTTTTGCGGAACAATCAAATATATTT | 48 | POLY |
| 36[34] | 36[6] | TTGGCAAATCAACAGTTGACCTAATTTACGA | 31 | POLY |
| 37[6] | 38[6] | ATAATATCCCATAAGGAATTGAGGACAAACAATTCGATCGCCATATTT | 48 | POLY |
| 35[0] | 34[21] | CTTATCCGGAAAGCATCACCTTGCTAGCAAAT | 32 | POLY |
| 33[1] | 34[0] | TATTATTTACTGATAGCCGAAAAATCTTATTCTAAG | 36 | POLY |

| Start | End | Sequence | Bases | ID |
|---|---|---|---|---|
| 7[14] | 8[6] | GGTGCCATCCCACGCAGTTCTAGCTGATCATTAAATTTTTGTTAAATTAAACGATG | 56 | POLY |
| 29[0] | 30[0] | TACGCAGTATGTAAGCGAAAGGAGACCCGCCGCAAGCCCAAT | 42 | POLY |
| 31[0] | 32[0] | AATTGAGTTGCTTAATGCATGCGCGAATCCCAATCC | 36 | POLY |
| 27[13] | 26[13] | CCATAGTGGTTCCGAAATCGGAAAATCCTGTTTGATGAATCA | 42 | POLY |
| 3[13] | 2[14] | CTGCCCGCTTTCCAGTGCTGACGCGACGGGAAACCTGTCGTG | 42 | POLY |
| 25[19] | 24[19] | CCTTTTGATAAGCAGTGTCACTGCGCGCATCAGACGATCCAGCGAGGTCATTTTTG | 56 | POLY |
| 23[19] | 22[20] | CTGTTTAGCTATTGCCGGGTTACCTGCACGTTAACGGCATCAGAATTTTCATTTGG | 56 | POLY |
| 19[14] | 18[14] | TGAACGGTAGATCACAGCGATCGTAAAA | 28 | POLY |
| 17[13] | 16[14] | TGTTAAATCAACGGACGGGAGCTCATTT | 28 | POLY |
| 21[20] | 20[20] | TATATTTTAAATATCGACATAAAAAATTTGGGCGGTTGTGTACGCAATGCCTGAG | 56 | POLY |
| 15[13] | 17[34] | TCGTAACCGGGGTTTTCCCAGTCACGACGAGGTGGAGCCGCCATAACCT | 49 | POLY |
| 11[6] | 12[6] | TTGTGAGAGAATGGGATAGGTCCGGAAACCAGGTGTGCTGCA | 42 | POLY |

| Start | End | Sequence | Bases | ID |
|---|---|---|---|---|
| 7[14] | 8[6] | GGTGCCATCCCACGCAGTTCTAGCTGATCATTAAATTTTTGTTAAATTAAACGATG | 56 | POLY |
| 13[6] | 14[14] | AAAGGGGGACAAAGCGCCATTCGCCATTTGGGTAACGCCATGCATCTGC | 49 | POLY |
| 1[13] | 0[13] | ATCCTGTTTGATATTTAGTTTGACCATTTTCTGCGAACGAGTAGGGTGGTTCCGAA | 56 | POLY |
| 7[14] | 8[6] | GGTGCCATCCCACGCAGTTCTAGCTGATCATTAAATTTTTGTTAAATTAAACGATG | 56 | POLY |
| 7[42] | 6[14] | TGCCGGAAATCACCATCAATATGATATTCAACCACCAGCTTACGGCTGG | 49 | POLY |
| 9[6] | 10[6] | GGGTAAAGTCAGCTCATTTTTCGGATTGACCGTATAGACTTT | 42 | POLY |
| 55[2] | 42[2] | GCGAGAAAGGAATGATAAATAAGGATATAATCCAGAGTCAAT | 42 | POLY |
| 53[6] | 54[3] | ACTATCGGCCTTGCTGGTACTCCGGCTTAGGGTTTGAAATACCGACCGTGGGGAAGAAAGCG | 62 | POLY |
| 51[6] | 52[6] | GAATACGTGGCAAGCTTAGATTAATACCTTTTTAACATATCCAGAACA | 48 | POLY |
| 5[14] | 4[13] | TCAGCAAATCGTTAACTCAACTTATTGGCATCAGATGCCGGG | 42 | POLY |
| 49[6] | 50[6] | AATCCTTTGCCCGAATTACCTTTTAAACATAGCGATCAGACAATATTT | 48 | POLY |
| 48[34] | 48[6] | ACATTTAACAATTTCATTTGAACGTTATTAA | 31 | POLY |

| Start | End | Sequence | Bases | ID |
|---|---|---|---|---|
| 7[14] | 8[6] | GGTGCCATCCCACGCAGTTCTAGCTGATCATTAAATTTTTGTTAAATTAAACGATG | 56 | POLY |
| 45[2] | 46[2] | ATGAAACAACAGATGAATATTCATTTCAGAAATAAA | 36 | POLY |
| 47[2] | 46[21] | ACGTAAAACAATTACCTGAGCAAAAGCGAATT | 32 | POLY |
| 43[2] | 44[2] | ACGCTGAGATGATTGTTTGTTTAACGTACATCAAGA | 36 | POLY |
| 41[7] | 28[0] | GAGAAAACTTTTAGAAACCACCAGGCGAGAAAGGAAGGGAAGATAGCAAACGTAG | 55 | POLY |
| 39[7] | 40[8] | GCTTAATTGAGAACAACTCGTATTTTTGCGGAACAATCAAATATATTT | 48 | POLY |
| 36[34] | 36[6] | TTGGCAAATCAACAGTTGACCTAATTTACGA | 31 | POLY |
| 37[6] | 38[6] | ATAATATCCCATAAGGAATTGAGGACAAACAATTCGATCGCCATATTT | 48 | POLY |
| 35[0] | 34[21] | CTTATCCGGAAAGCATCACCTTGCTAGCAAAT | 32 | POLY |
| 33[1] | 34[0] | TATTATTTACTGATAGCCGAAAAATCTTATTCTAAG | 36 | POLY |
| 29[0] | 30[0] | TACGCAGTATGTAAGCGAAAGGAGACCCGCCGCAAGCCCAAT | 42 | POLY |
| 31[0] | 32[0] | AATTGAGTTGCTTAATGCATGCGCGAATCCCAATCC | 36 | POLY |
| 27[13] | 26[13] | CCATAGTGGTTCCGAAATCGGAAAATCCTGTTTGATGAATCA | 42 | POLY |

| Start | End | Sequence | Bases | ID |
| --- | --- | --- | --- | --- |
| 7[14] | 8[6] | GGTGCCATCCCACGCAGTTCTAGCTGATCATTAAATTTTTGTTAAATTAAACGATG | 56 | POLY |
| 3[13] | 2[14] | CTGCCCGCTTTCCAGTGCTGACGCGACGGGAAACCTGTCGTG | 42 | POLY |
| 25[19] | 24[19] | CCTTTTGATAAGCAGTGTCACTGCGCGCATCAGACGATCCAGCGAGGTCATTTTTG | 56 | POLY |
| 23[19] | 22[20] | CTGTTTAGCTATTGCCGGGTTACCTGCACGTTAACGGCATCAGAATTTTCATTTGG | 56 | POLY |
| 19[14] | 18[14] | TGAACGGTAGATCACAGCGATCGTAAAA | 28 | POLY |
| 17[13] | 16[14] | TGTTAAATCAACGGACGGGAGCTCATTT | 28 | POLY |
| 21[20] | 20[20] | TATATTTTAAATATCGACATAAAAAAATTTGGGCGGTTGTGTACGCAATGCCTGAG | 56 | POLY |
| 15[13] | 17[34] | TCGTAACCGGGGTTTCCCAGTCACGACGAGGTGGAGCCGCCATAACCT | 49 | POLY |
| 11[6] | 12[6] | TTGTGAGAGAATGGGATAGGTCCGGAAACCAGGTGTGCTGCA | 42 | POLY |
| 13[6] | 14[14] | AAAGGGGGACAAAGCGCCATTCGCCATTTGGGTAACGCCATGCATCTGC | 49 | POLY |
| 1[13] | 0[13] | ATCCTGTTTGATATTTAGTTTGACCATTTTCTGCGAACGAGTAGGGTGGTTCCGAA | 56 | POLY |

**Table S4A.** List of Padlock Strands Sequences (H1)
This table denotes the ssDNA padlocking sequences used for the H1 hinge. The Start and End columns denote the helix[base] positions at the 5' and 3' ends of the DNA staple strands in the CaDNAno design for the respective designs (Supplementary Fig. 2A). Refer to Table 1 for the definitions and colors of the IDs.

| Start | End | Sequence | Bases | ID |
|---|---|---|---|---|
| 20[76] | 34[63] | TATCATCCATGCATTTTTTTTTTTTTTT<br>CAATATAATATAGTATAAGATTTTTTTT<br>TTTTTTCATGCATTGTAGAACGGTATG | 83 | PADLOCK1 |
| 25[63] | 29[76] | GCAAATTGCCGCGCCATGCATTTTTTT<br>TTTTTTTTCAATATAATATAGTATAAGA<br>TTTTTTTTTTTTTTCATGCATCACCGCC | 83 | PADLOCK2 |

**Table S4B.** List of Padlocking Strands Sequences (H2)
This table denotes the ssDNA padlocking sequences used for the H2 hinge. The Start and End columns denote the helix[base] positions at the 5' and 3' ends of the DNA staple strands in the CaDNAno design for the respective designs (Supplementary Fig. 2B). Refer to Table 1 for the definitions and colors of the IDs.

| Start | End | Sequence | Bases | ID |
|---|---|---|---|---|
| 25[63] | 29[76] | TTTCACGGCCTGGCCATGCATTTTTTT<br>TTTTTTTTCAATATAATATAGTATAAGA<br>TTTTTTTTTTTTTTCATGCATCGCTTTC<br>TAAAGCA | 90 | PADLOCK1 |
| 19[91] | 34[84] | AAGGAACTGTCGTCCATGCATTTTTTT<br>TTTTTTTTCAATATAATATAGTATAAGA<br>TTTTTTTTTTTTTTCATGCATGCCACC<br>ATGCCAAC | 90 | PADLOCK2 |

**Table S5 .** List of Locking Mechanism Strands
This table outlines the ssDNA strands that can bind specifically to the padlock strands.

| Sequence | Bases | ID |
|---|---|---|
| AAAAAAAAAAAAAAAAAACGGCTT | 24 | LOCK |
| AGCCGTTTTTTTTTTTTTTTTTT | 23 | ANTI-LOCK |
| CGGCTTAAAAAAATCTTATACTATATT ATATTCAAAAAAA | 40 | KEY |
| TTTTTTTCAATATAATATAGTATAAGAT TTTTTTAAGCCG | 40 | ANTI-KEY |

### Supporting Files

Scripts and .json files can be found at:
https://github.com/ubcbiomod/Higher-Order-Nanohinge-Systems
